# Extracellular ISG15 triggers ISGylation via a type-I interferon independent non-canonical mechanism to regulate host response during virus infection

**DOI:** 10.1101/2024.07.05.602290

**Authors:** Lindsay Grace Miller, Kim Chiok, Charles Mariasoosai, Indira Mohanty, Sudiksha Pandit, Pallavi Deol, Liyon Mehari, Michael N. Teng, Arthur L. Haas, Senthil Natesan, Tanya A. Miura, Santanu Bose

## Abstract

Type-I interferons (IFN) induce cellular proteins with antiviral activity. One such protein is Interferon Stimulated Gene 15 (ISG15). ISG15 is conjugated to proteins during ISGylation to confer antiviral activity and regulate cellular activities associated with inflammatory and neurodegenerative diseases and cancer. Apart from ISGylation, unconjugated free ISG15 is also released from cells during various conditions, including virus infection. The role of extracellular ISG15 during virus infection was unknown. We show that extracellular ISG15 triggers ISGylation and acts as a soluble antiviral factor to restrict virus infection via an IFN-independent mechanism. Specifically, extracellular ISG15 acts post-translationally to markedly enhance the stability of basal intracellular ISG15 protein levels to support ISGylation. Furthermore, extracellular ISG15 interacts with cell surface integrin (α5β1 integrins) molecules via its RGD-like motif to activate the integrin-FAK (Focal Adhesion Kinase) pathway resulting in IFN-independent ISGylation. Thus, our studies have identified extracellular ISG15 protein as a new soluble antiviral factor that confers IFN-independent non-canonical ISGylation via the integrin-FAK pathway by post-translational stabilization of intracellular ISG15 protein.

## Introduction

Type-I Interferons (IFN) mediate antiviral responses via induction of effectors known as Interferon-Stimulated Genes (ISGs)^1,2^. Interferon Stimulated Gene 15 (ISG15) is an ISG involved in antiviral response and immunomodulatory activity^3–9^. ISG15 is post-translationally conjugated to host and viral proteins by three ubiquitin-like conjugating enzymes (UBE1L, UBCH8 and HERC5) in a process known as ISGylation^10–13^. ISGylation is a key biological activity that regulates various diseases including infectious diseases, cancer, and neurodegenerative disorders^14^. More importantly, ISGylation confers potent wide-spectrum antiviral activity against large group of RNA and DNA viruses including human respiratory syncytial virus (RSV), Ebola virus, influenza A virus (IAV), rabies virus, Zika virus, dengue virus, West Nile virus (WNV), lymphocytic choriomeningitis virus (LCMV), human immunodeficiency virus-1 (HIV-1), hepatitis B virus (HBV), hepatitis C virus (HCV), coxsackievirus B3 (CVB3), Sindbis virus, Kaposi’s sarcoma-associated herpesvirus (KSHV), herpes simplex type-1 (HSV-1), and cytomegalovirus (CMV)^15–26^. Although ISGylation occurs in the cytoplasm, several studies have observed the release of unconjugated monomeric ISG15 protein into the extracellular milieu^5,27,28^. Soluble, extracellular ISG15 promotes proliferation of and exerts cytokine-like activity in immune cells (natural killer cells, PBMCs, T cells) that culminates in IFN-γ production^5,28^.

IFNs play an important role in the innate immune antiviral response against various viruses including respiratory viruses^29–34^. Although viral (SARS-CoV-2^35,36^, IAV^36^, and rhinovirus^37^) infection triggers extracellular release of ISG15 from infected cells, the role of extracellular ISG15 during virus infection is not well understood. Furthermore, the virus-associated mechanisms by which infection triggers the release of ISG15 and whether extracellular ISG15 has direct antiviral activity during virus infection remain unknown. Although ISG15 is an IFN-stimulated gene^3–13^, our current study surprisingly revealed that extracellular ISG15 induces an increase in intracellular ISG15 protein levels during ISGylation via an IFN-independent mechanism. Intriguingly, by using two clinically relevant respiratory RNA viruses, RSV and SARS-CoV-2, we show that extracellular ISG15 possesses antiviral activity and can restrict virus infection independent of IFN. Mechanistically, extracellular ISG15 interacted with the cell surface α5β1 integrin to enhance intracellular ISG15 protein levels and ISGylation in target cells via an integrin-Focal Adhesion kinase (FAK) pathway. Extracellular ISG15 utilized an IFN-independent mechanism for regulating intracellular ISG15 levels post-translationally by enhancing the stability of intracellular ISG15 protein. We also identified the envelope proteins of SARS-CoV-2 (Spike or S protein) and RSV (Fusion or F protein) as key viral factors involved in stimulating ISGylation and ISG15 release. Thus, we have identified extracellular ISG15 as a new soluble antiviral factor and mechanistic studies unfolded a novel IFN-independent antiviral response by extracellular ISG15, which is mediated by integrin-FAK signaling. Furthermore, extracellular ISG15 regulated intracellular ISG15 levels independent of IFN by enhancing the stability of ISG15 protein during ISGylation. ISGylation is an important cellular activity, and our study has highlighted a new non-canonical mechanism triggering ISGylation, which may have wider ramifications for controlling a wide spectrum of both viral and non-infectious diseases.

## Results

### SARS-CoV-2 S protein induces intracellular ISG15, ISGylation, and extracellular release of ISG15 in epithelial cells

SARS-CoV-2 inhibits ISGylation response in infected myeloid immune cells (macrophages) due to the activity of the virally encoded ISGylation antagonist, PLpro. The PLpro removes conjugated ISG15 from its cellular targets^35,38^. In infected macrophages, the PLpro de-ISGylase activity leads to loss of ISGylation and release of ISG15 into the extracellular milieu^36^. However, whether non-myeloid epithelial cells infected with SARS-CoV-2 undergo ISGylation and release of ISG15 is unknown. Therefore, we aimed to determine whether infection with SARS-CoV-2 resulted in ISGylation and extracellular release of ISG15 from epithelial cells. We infected human lung epithelial Calu-3 cells with SARS-CoV-2 and examined ISGylation-ISG15 dynamics by immunoblotting with ISG15 antibody (Fig. 1A). As expected, ISGylation was not evident at 48h post-infection, though unconjugated ISG15 was increased. Remarkably, ISGylation was readily detected in SARS-CoV-2 infected cells at 120h post-infection (Fig. 1A). ISGylation status in infected cells correlated with PLpro expression, since in contrast to 48h post-infection, PLpro was undetectable at 120h post-infection (Fig. 1A). Immunoblot analysis of TCA precipitated proteins from medium supernatant revealed SARS-CoV-2 infection stimulating extracellular release of ISG15 from infected Calu-3 cells at 120h post-infection (Fig. 1B). This ISG15 release corresponded with 37% cell death in infected cells at 120h post-infection (Fig. 1C). These results indicate that SARS-CoV-2 infection of human lung epithelial cells induced ISGylation and extracellular release of ISG15 late during infection.

**Figure 1.**
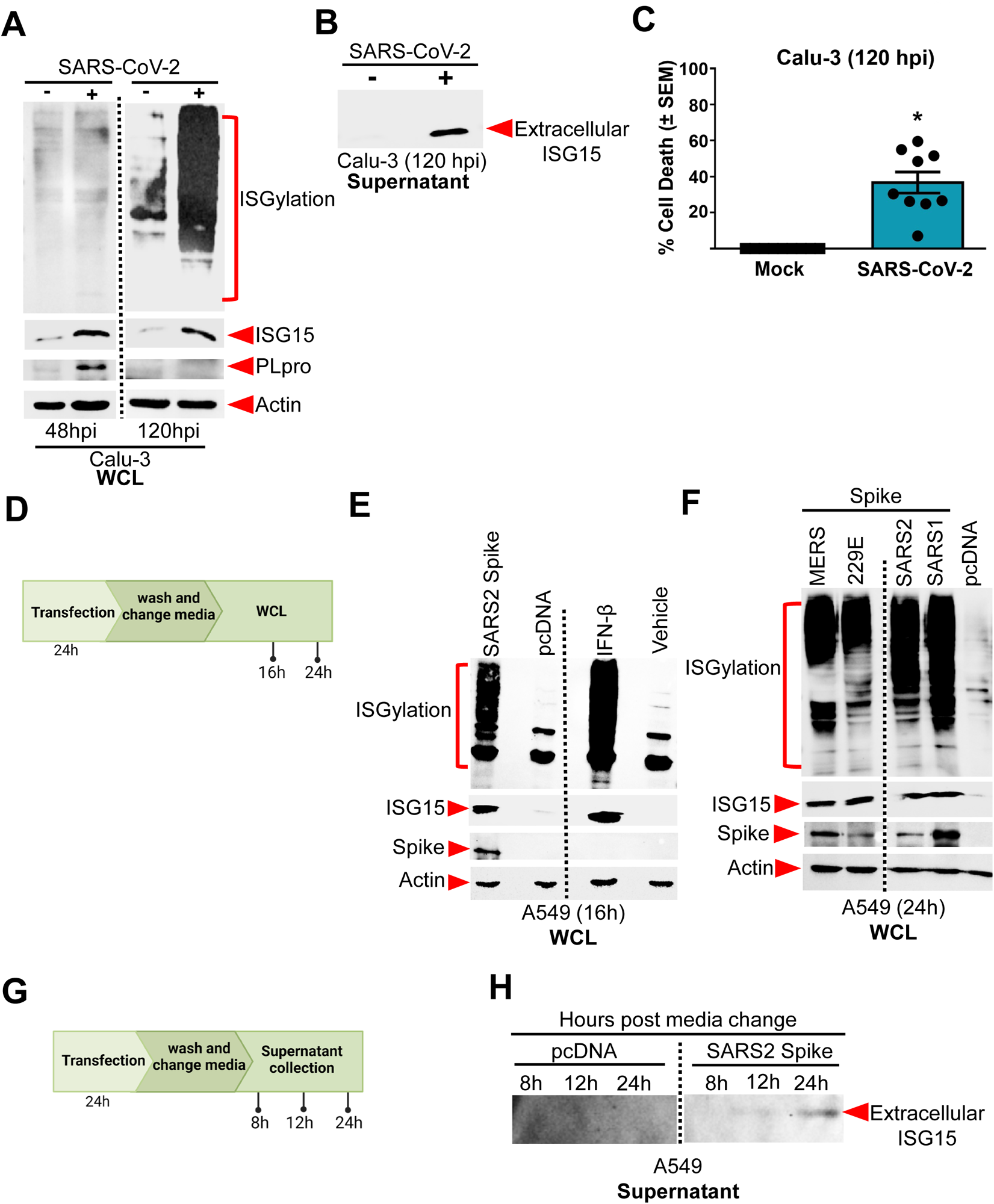
SARS-CoV-2 S protein induces expression of intracellular ISG15, ISGylation, and extracellular release of ISG15 in epithelial cells. A) Human lung epithelial Calu-3 cells were infected with SARS-CoV-2 (MOI= 0.1). Whole cell lysate (WCL) collected at 48h post-infection (hpi) and 120hpi was subjected to immunoblotting with ISG15, PLpro and actin antibodies. B) Cell culture supernatants (supernatant) from SARS-CoV-2 (MOI= 0.1) infected Calu-3 cells were collected at 120h post infection. Trichloroacetic acid (TCA) precipitated supernatant was subjected to immunoblot with anti-ISG15 antibody to detect extracellular ISG15. C) Supernatants from mock and SARS-CoV-2 infected (120 hpi) Calu-3 cells were used to quantify cell death by lactate dehydrogenase (LDH) assay. D) Experimental schematic – human lung epithelial A549 cells were transfected with either plasmids encoding C9-tagged S proteins from various coronaviruses or empty vector control plasmid pcDNA 3.1 (pcDNA). At 24h post-transfection, washed cells were incubated with fresh media without plasmids for 16h and 24h. Whole cell lysate (WCL) was collected for analysis. E) WCL collected from A549 cells transfected with C9-tagged SARS-CoV-2 (SARS2) S protein as described in the experimental schematic in Fig. 1D was subjected to immunoblotting with ISG15, C9, and actin antibodies. A549 cells treated with interferon-β (IFN-β, 100U/mL) for 24 was used as a positive control for ISGylation. F) A549 cells were transfected with C9-tagged S protein from various coronaviruses including MERS-CoV (MERS), 229E, SARS-CoV-2 (SARS2) and SARS-CoV-1 (SARS1) or empty vector control (pcDNA). The experiment was conducted as described in the experimental scheme shown in Fig. 1D. WCL collected at 24h was subjected to immunoblotting with ISG15, C9, and actin antibodies. G) Experimental schematic - A549 cells were transfected with either plasmid encoding C9-tagged SARS-CoV-2 S protein or empty vector control (pcDNA). At 24h post-transfection, washed cells were incubated with fresh media without plasmids for 8h, 16h and 24h. Total protein in plasmid-free culture supernatant was precipitated with TCA. H) TCA precipitated culture supernatant collected from A549 cells transfected with C9-tagged SARS-CoV-2 (SARS2) S protein as described in the experimental schematic in Fig. 1G was subjected to immunoblotting with ISG15 antibody to detect extracellularly released ISG15. The immunoblots are representative of data from three independent experiments with similar results.

Since ISGylation and ISG15 release occurred in SARS-CoV-2 infected epithelial cells (Fig. 1A, 1B), we next examined whether viral encoded factor(s) could be responsible for such activity. We focused on the role of the SARS-CoV-2 spike (S) protein because previous reports have shown upregulation of ISG15 in cells expressing the S protein^39^. Nevertheless, it is still unknown whether SARS-CoV-2 S protein can trigger ISGylation and ISG15 release. We expressed human coronavirus S protein tagged with a C9 peptide in human lung epithelial A549 cells. A549 cells were transfected with the plasmids for 24h and were then washed and incubated with plasmid-free medium for an additional 16h and 24h prior to collection of cell lysates for immunoblotting (Fig. 1D). In contrast to empty control plasmid, ectopic expression of the SARS-CoV-2 (SARS2) S protein promoted expression of intracellular ISG15 and ISGylation in A549 cells (Fig. 1E, left panel). As a positive control, we treated A549 cells with IFN-β, a potent ISGylation inducer (Fig. 1E, right panel). Expression of S proteins from the human coronaviruses SARS-CoV-1 (SARS1), Middle East Respiratory Syndrome coronavirus (MERS) and 229E also led to enhanced intracellular ISG15 expression and ISGylation in A549 cells (Fig. 1F), suggesting that intracellular expression of S proteins from various coronaviruses can trigger ISGylation. Extracellular SARS-CoV-2 protein can also trigger various cellular responses, including pro-inflammatory response^40^. Therefore, we examined whether extracellular S protein can trigger ISGylation. In contrast to intracellular expression of S protein, A549 cells treated with purified SARS-CoV-2 S protein (full length or the S1 subunit) failed to induce ISG15 expression or ISGylation (Supp. Fig. 1A). Thus, S protein expression in coronavirus infected epithelial cells may represent a common mechanism driving ISGylation, which we show occurring during late infection of SARS-CoV-2 infected epithelial cells, after PLpro was no longer detected (Fig. 1A). We next evaluated whether the S protein can also promote extracellular release of unconjugated ISG15. Cells expressing the SARS-CoV-2 (SARS2) S protein released ISG15 into the medium as shown by immunoblotting of TCA precipitated medium supernatant (Fig. 1H). Thus, the S protein of SARS-CoV-2 is a viral factor that not only triggers expression of ISG15 and ISGylation, but it also facilitates extracellular release of unconjugated ISG15.

### Infection with Respiratory Syncytial virus (RSV) induces ISGylation and extracellular release of ISG15 from epithelial cells: RSV Fusion (F) protein triggers ISGylation and ISG15 release

We next investigated whether infection with RSV, a respiratory RNA virus that lacks de-ISGylases (e.g, PLpro of SARS-CoV-2) promotes ISGylation and release of ISG15 from epithelial cells. We infected human lung epithelial A549 cells with recombinant RSV expressing mKate2 red fluorescent protein (mKate2-RSV)^41^. We used RFP (red fluorescent protein) antibody to monitor mKate2-RSV infection status by immunoblotting. RSV infection induced intracellular ISG15 protein and ISGylation within 48h post-infection (Fig. 2A). We also detected extracellular release of ISG15 at 48h post-infection (Fig. 2B). In contrast to SARS-CoV-2 (Fig. 1C), ISG15 release from RSV infected A549 cells occurred independent of lytic cell death since cell death at 48h post-infection was negligible (i.e., less than 3% cell death) (Fig. 2C). Although ISGylation has been noted in RSV infected cells by us (Fig. 2A) and other studies^42,43^, ISG15 release from RSV infected cells had not been reported. Our results show that RSV is another respiratory RNA virus that mediates the extracellular release of ISG15 during infection.

**Figure 2.**
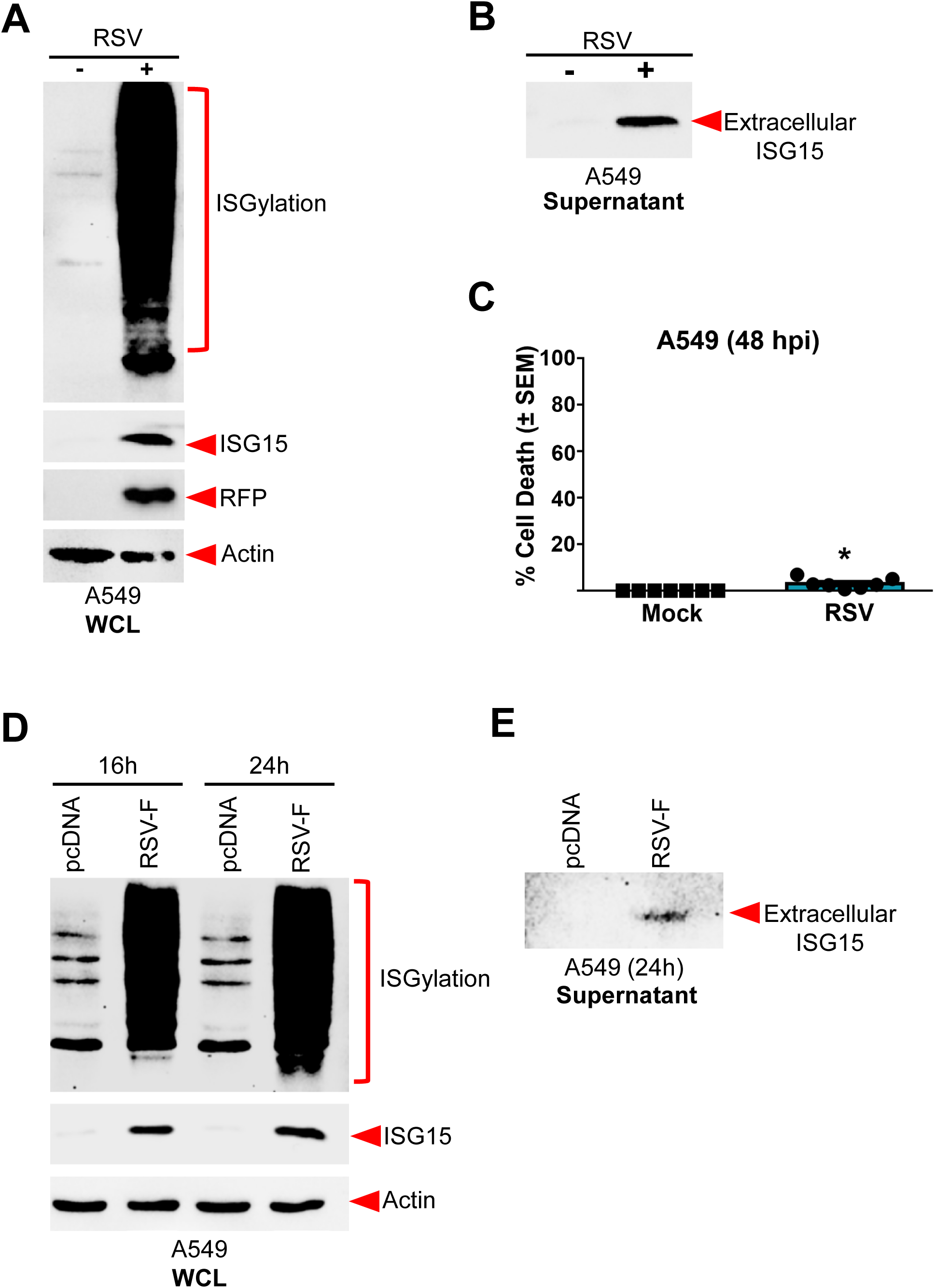
Respiratory syncytial virus (RSV) induces ISGylation and extracellular release of ISG15 from epithelial cells: RSV Fusion (F) protein triggers ISGylation and ISG15 release. A) Human lung epithelial A549 cells were infected with RSV expressing mKate2 protein (MOI=0.1). WCL collected at 48h post-infection was subjected to immunoblotting with ISG15, RFP (to detect mKate2 protein) and actin antibodies. B) Cell culture supernatants from A549 cells infected with RSV as described in Fig. 2A were precipitated with TCA and total protein was immunoblotted with ISG15 antibody to detect extracellular ISG15. C) Supernatants from mock and RSV infected (48h post-infection) A549 cells were used to quantify cell death by lactate dehydrogenase (LDH) assay. D) A549 cells were transfected with either empty vector control plasmid (pcDNA) or RSV F protein plasmid. At 24h post-transfection, washed cells were incubated with fresh media without plasmids for 16h and 24h. Whole cell lysate (WCL) was subjected to immunoblotting with ISG15 and actin antibodies. E) A549 cells were transfected with either empty vector control plasmid (pcDNA) or RSV F protein plasmid. At 24h post-transfection, washed cells were incubated with fresh media without plasmids for 24h. Total protein in plasmid-free culture supernatant was precipitated with TCA. TCA precipitated culture supernatant was subjected to immunoblotting with ISG15 antibody to detect extracellularly released ISG15. The immunoblots are representative of data from three independent experiments with similar results. \**p* < 0.05 using a Student’s t-test.

Since we identified the envelope S protein of SARS-CoV-2 as a viral factor promoting ISGylation and ISG15 release, we evaluated similar activity of envelope Fusion (F) protein of RSV. For these studies, we expressed RSV F protein in A549 cells. As with the S protein experiment, we transfected A549 cells with either pcDNA (control) or RSV F plasmids. At 24h post-transfection, cells were washed and incubated with plasmid-free medium for an additional 24h. Immunoblotting of the cell lysate with RSV F antibody revealed efficient expression of F in transfected cells (Supp. Fig. 1B). We repeated the experiment by collecting cell lysate either 16h or 24h post-incubation with plasmid free medium. Immunoblotting demonstrated robust induction of intracellular ISG15 protein and ISGylation by RSV F protein (Fig. 2D). Expression of RSV F protein also resulted in release of ISG15 from A549 cells as deduced by immunoblotting of TCA precipitated medium supernatant with ISG15 antibody (Fig. 2E). These results have identified the RSV F protein as a viral encoded factor promoting ISGylation and extracellular release of ISG15. Thus, our studies have identified viral envelope proteins of two respiratory RNA viruses (i.e., S protein of SARS-CoV-2 and F protein of RSV) in triggering ISGylation and ISG15 release.

### Extracellular ISG15 induces intracellular ISG15 and ISGylation

Two clinically important human pathogens, SARS-CoV-2 and RSV triggered extracellular release of ISG15 from lung epithelial cells during infection (Fig. 1B, 1H, 2B, 2E). To dissect the role of extracellular ISG15, we used purified recombinant ISG15 protein (rISG15) (Supp. Fig. 2A) to investigate whether extracellular ISG15 can induce intracellular ISG15 protein and ISGylation in uninfected cells. Treatment of two human lung epithelial cell lines, A549 (Fig. 3A) and Calu-3 (Fig. 3B), with purified rISG15 led to induction of intracellular ISG15 protein levels and ISGylation in both cell types as early as 6h post-treatment as deduced by immunoblotting. ISG15 detected in the blot constitutes intracellular ISG15 protein since it was detected after trypsin treatment (to remove extracellular rISG15) of rISG15 incubated cells (Supp. Fig. 2B, 2C). Additionally, we failed to detect extracellularly added (and incubated for 6h) biotinylated rISG15 (biot-ISG15) in trypsin treated A549 cell lysate following immunoblotting with avidin antibody (Supp. Fig. 2D). This result demonstrated that extracellular ISG15 is not internalized.

**Figure 3.**
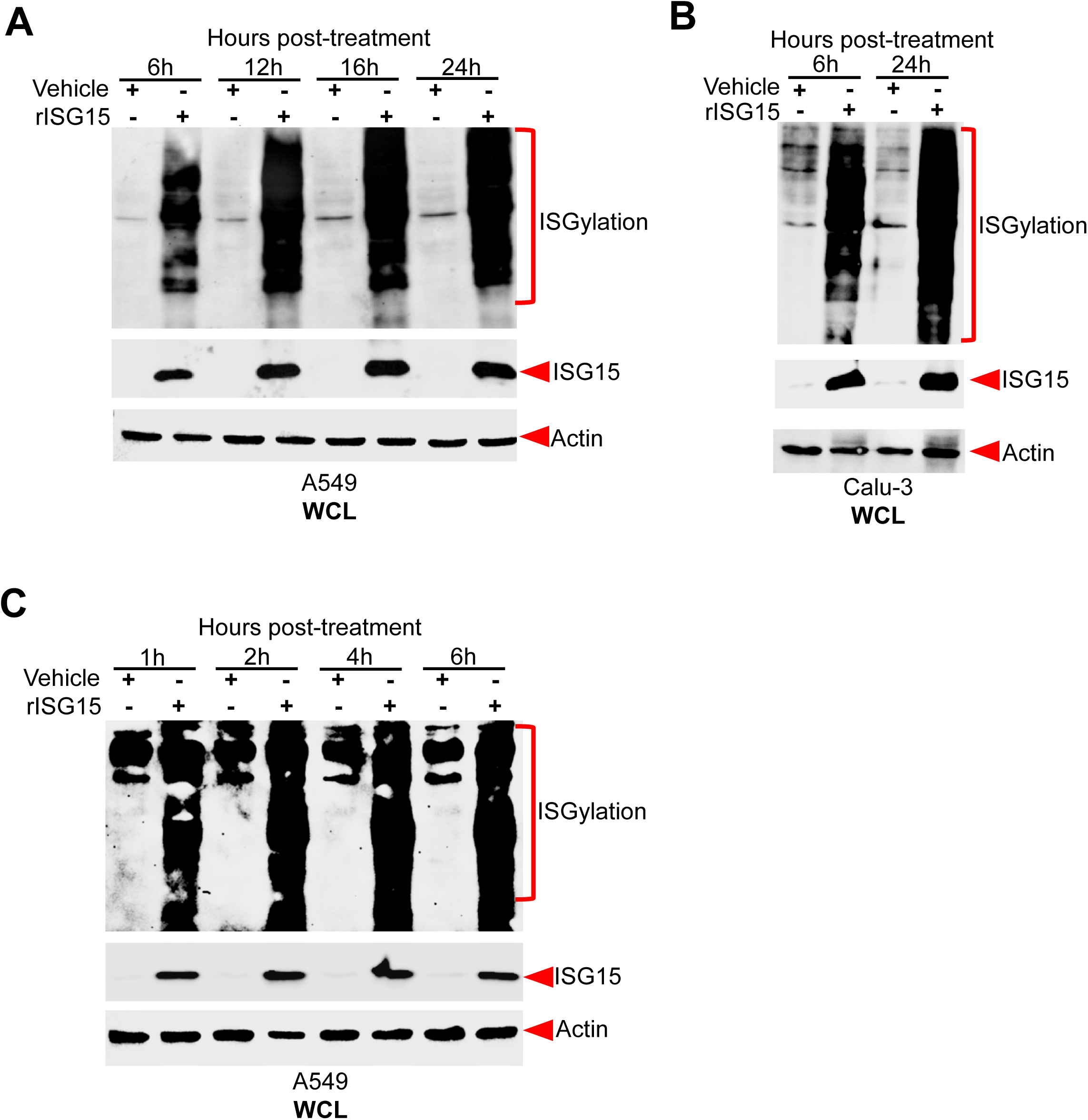
Extracellular ISG15 induces intracellular ISG15 and ISGylation. Human lung epithelial A549 (A) and Calu-3 (B) cells were treated with either vehicle control or purified recombinant ISG15 protein (rISG15, 5µg/mL) for the indicated times and WCL was immunoblotted with ISG15 and actin antibodies. C) A549 cells were treated with rISG15 for indicated time-period. WCL was immunoblotted with ISG15 and actin antibodies. The immunoblots are representative of data from three independent experiments with similar results.

ISGylation at 6h post-treatment with rISG15 (Fig. 3A, 3B) suggests that extracellular ISG15 may rapidly enhance the pool of intracellular ISG15 protein levels for ISGylation. Therefore, we investigated the kinetics of intracellular ISG15 protein induction and ISGylation in rISG15 treated A549 cells. Extracellular ISG15 rapidly induced intracellular ISG15 protein and ISGylation in A549 cells as early as 1h post-treatment with rISG15 (Fig. 3C). While extracellular ISG15 has a cytokine-like activity in immune cells^5,27,44^, our study has identified a previously unknown pro-ISGylation activity of extracellular ISG15.

### Extracellular ISG15 induces intracellular ISG15 protein and ISGylation in target cells via a type-I interferon-independent mechanism

Since purified ISG15 protein (rISG15) rapidly and robustly induced intracellular ISG15 protein and ISGylation (Fig. 3), we investigated whether the ISGylation activity of extracellular ISG15 is mediated via an IFN-dependent mechanism.

Wild-type 2fTGH cells and their counterpart U5A cells, which lack a functional type-I interferon (IFN) receptor (IFNAR), were used to probe the involvement of IFN during extracellular ISG15 mediated response. These cells also lack type-III IFN signaling (IFN-λ)^45,46^. Additionally, type-II (IFN-γ) signaling is lacking in 2fTGH and U5A cells since they are non-immune cells. As expected, in contrast to wild-type 2fTGH control cells, U5A cells lacked ISGylation and expression of intracellular ISG15 following treatment with IFN-β (Supp. Fig. 3A). In contrast, both 2fTGH and U5A cells displayed robust induction of intracellular ISG15 protein and ISGylation following treatment with purified ISG15 protein (rISG15) (Fig. 4A). This unexpected result was validated using VeroE6 epithelial cells which are unable to produce IFN. Like U5A cells, treatment of VeroE6 cells with rISG15 led to rapid and strong induction of intracellular ISG15 and ISGylation (Fig. 4B). These results together demonstrate that extracellular ISG15-mediated induction of intracellular ISG15 protein and ISGylation occurs through an IFN-independent mechanism.

**Figure 4.**
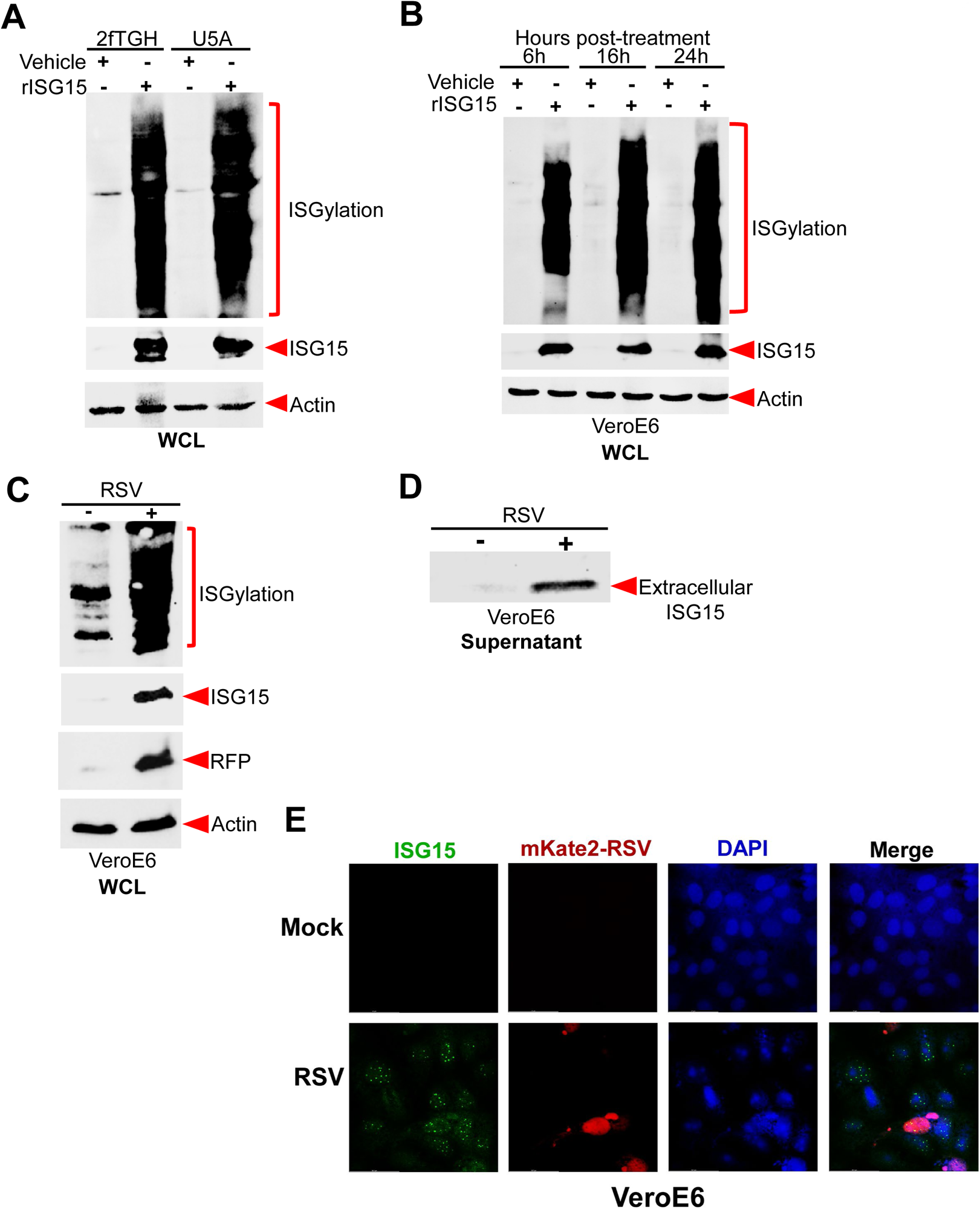
Extracellular ISG15 induces intracellular ISG15 protein and ISGylation via a type-I interferon (IFN) independent mechanism. A) 2fTGH (IFN-competent) and U5A (IFN-incompetent) cells were treated with purified recombinant ISG15 protein (rISG15, 5µg/mL) for 6h before collection of WCL for immunoblotting with ISG15 and actin antibodies. B) VeroE6 cells (IFN-incompetent) were treated with rISG15 (5µg/mL) for 6h, 16h and 24h before collection of WCL for immunoblotting with ISG15 and actin antibodies. C) VeroE6 cells were infected with mKate2-RSV (MOI=0.1) for 48h before WCL collection for immunoblotting with ISG15, RFP (to detect mKate2) and actin antibodies. D) Cell culture supernatants from VeroE6 cells infected with RSV as described in Fig. 4C were precipitated with TCA and total protein was immunoblotted with ISG15 antibody to detect extracellular ISG15. E) Confocal microscopic image of VeoE6 cells mock-infected or infected with mKate2-RSV (MOI=0.1). Cells were labelled with ISG15 antibody and CoraLite488-secondary antibody (green) to detect endogenous ISG15 protein. Nuclei were stained with DAPI (blue). Infected cells were visualized by the red fluorescence emitted from the mKate2 protein. The immunoblots are representative of data from three independent experiments with similar results.

Next, we evaluated whether viral infection could induce intracellular ISG15 protein and ISGylation in IFN-deficient VeroE6 cells. RSV infection robustly induced intracellular ISG15 protein and ISGylation in VeroE6 cells (Fig. 4C). Additionally, RSV infected VeroE6 cells were competent in extracellular release of ISG15 (Fig. 4D). As with infected A549 cells (Fig. 2C), the release of ISG15 from infected VeroE6 cells was independent of lytic cell death since cell death was not observed during the same timeframe as ISG15 release (Supp. Fig. 3B).

Virus infection resulted in the extracellular release of ISG15 (Fig. 1B, 2B, 4D) and extracellular ISG15 triggered intracellular ISG15 protein induction and ISGylation (Fig. 3A, 3B, 4A, 4B). This phenomenon occurring in IFN incompetent cells (VeroE6 and U5A) (Fig. 4A, B, C, D) suggested an IFN-independent mechanism. Thus, we postulate that ISG15 released from virus infected cells may act via paracrine action on uninfected bystander cells to promote ISG15 induction in those cells via IFN independent activity. To mimic this scenario, we infected IFN-lacking VeroE6 cells with mKate2-RSV at low MOI (MOI = 0.1) to ease the distinction between infected and uninfected cells. Fixed cells labelled with ISG15 antibody and a CoraLite488-labeled (green) secondary antibody were subjected to confocal fluorescence microscopy to detect infected cells (red cells - positive for mKate2) and cells expressing ISG15 (green cells). As expected, we visualized ISG15 in infected VeroE6 cells (Fig. 4E) (Supp. Fig. 3C) as evident from the merged images (yellow). Interestingly, we also visualized high levels of ISG15 in uninfected bystander cells (Fig. 4E) (Supp. Fig. 3C), suggesting an IFN-independent paracrine activity of extracellular ISG15 during virus infection. As intracellular ISG15 and ISGylation participate in innate antiviral defense^15–26^, we envision a paracrine antiviral function of extracellular ISG15 released from infected cells in priming or pre-activating the host defense apparatus in uninfected bystander cells prior to infection. Likewise, extracellular ISG15 may act on virus infected cells via an autocrine mechanism to amplify intracellular ISG15 levels and ISGylation in infected cells for optimal ISGylation-dependent antiviral response.

Thus, we have identified a novel activity of extracellular ISG15 in triggering induction of intracellular ISG15 protein and ISGylation via an IFN-independent mechanism. To the best of our knowledge, this is the first report of an IFN-independent mechanism for ISGylation during virus infection.

### IFN-independent antiviral activity of extracellular ISG15

Extracellular ISG15-mediated induction of intracellular ISG15 and ISGylation via an IFN-independent mechanism (Fig. 4) prompted us to explore its implications during virus infection. RSV infection triggered ISG15 release, and ISG15 protein was induced in bystander uninfected cells (Fig. 2 and Fig. 4). Therefore, we postulated that extracellular ISG15 may possess an innate host defense function that can limit virus infection.

To evaluate the antiviral activity of extracellular ISG15, we pre-treated human lung epithelial A549 cells with purified ISG15 protein (rISG15), followed by RSV infection. Extracellular ISG15 mediated restriction of RSV infectivity and spread was evident in phase-contrast microscopy showing reduced cytopathic effect (CPE) following rISG15 treatment versus extensive CPE in vehicle treated infected A549 cells (Supp. Fig. 4A). This result was confirmed by observing diminished virus replication in A549 cells treated with rISG15 and infected with mKate2-RSV (Supp. Fig. 4B). Most importantly, plaque assay analysis for infectious viral titer revealed loss of infectivity (reduction by one log or 10-fold) following extracellular rISG15 treatment of RSV infected A549 cells (Supp. Fig. 4C). Accordingly, a representative plaque assay for infectious viral titer also showed reduction in plaques from rISG15 treated infected A549 cells compared to control vehicle treated infected cells (Supp. Fig. 4D). These results demonstrated antiviral activity of soluble extracellular ISG15.

Identification of extracellular ISG15 as an antiviral factor prompted us to investigate whether such antiviral activity is mediated by the IFN response. For our studies, we used IFN-lacking VeroE6 cells. RSV infection in VeroE6 cells treated with rISG15 was analyzed using the same methods as described above for IFN-competent A549 cells (Supp. Fig. 4A, 4B, 4C, 4D). Although VeroE6 cells lack IFN, extracellular ISG15 still conferred protection against virus infection as inferred from - a) diminished viral replication as deduced from loss of mKate2 expression in mKate2-RSV infected VeroE6 cells treated with rISG15 (Fig. 5A), b) reduced virus-associated CPE in rISG15 treated cells (Fig. 5B), and c) plaque assays showing reduced infectious viral titer due to rISG15 treatment (Fig. 5C and Supp. Fig. 4E). Viral infectivity was significantly reduced by more than 10-fold following treatment of VeroE6 cells with rISG15 (Fig. 5C). Thus, we have identified extracellular ISG15 as a new soluble antiviral factor which operates via an IFN-independent mechanism.

**Figure 5.**
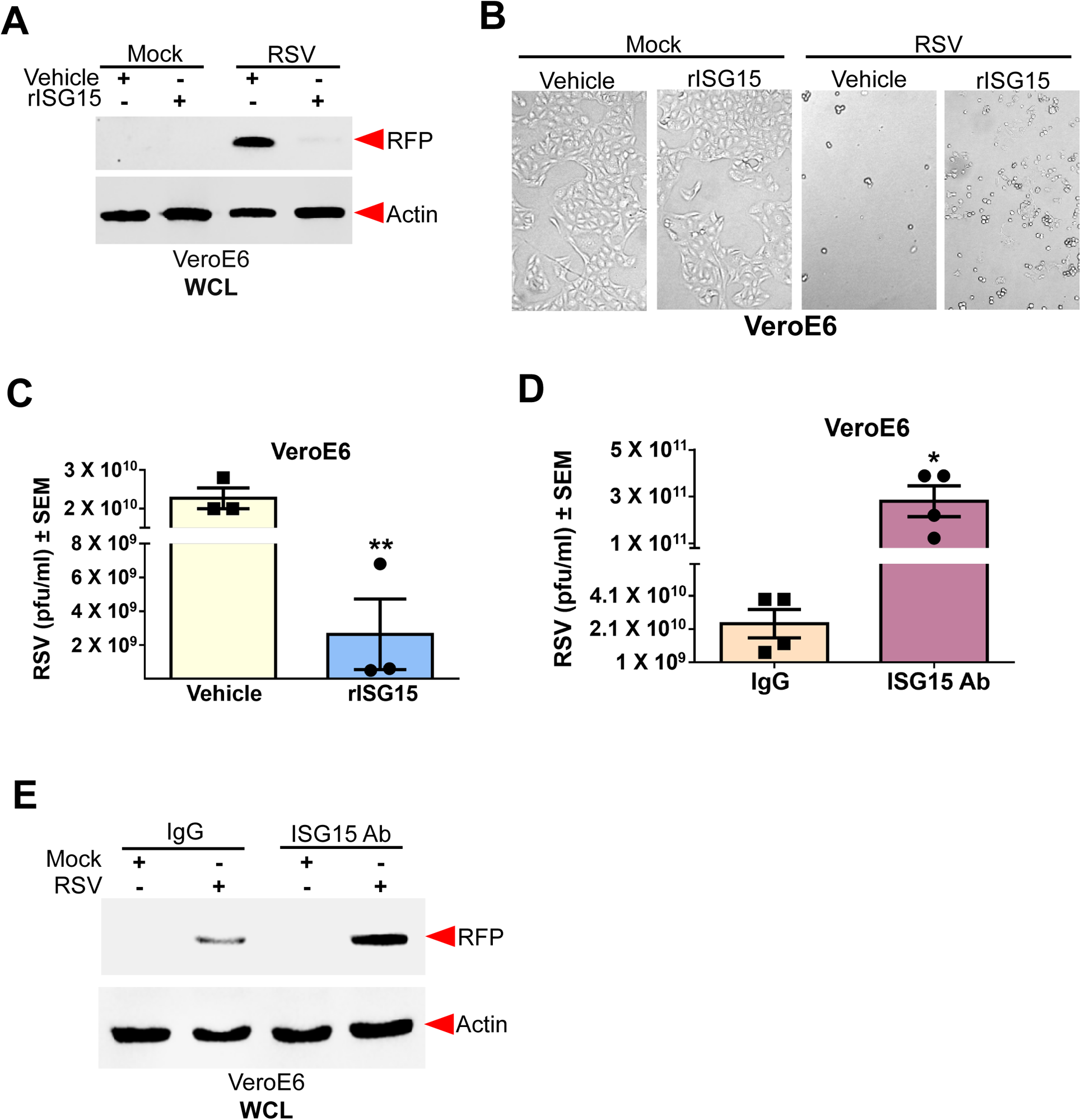
Antiviral activity of extracellular ISG15 protein: IFN-independent antiviral response by extracellular ISG15. A) VeroE6 cells were pre-treated with purified recombinant ISG15 protein (rISG15, 5µg/mL) or vehicle control for 8h prior to infection with mKate2-RSV for 48h before collection of WCL for immunoblotting with RFP (to detect mKate2) and actin antibodies. B) Bright field microscopy of VeroE6 monolayers treated with vehicle control or rISG15 prior to infection with RSV (MOI=1, 16h). C) Infectious viral titer following RSV (MOI=0.1) infection of VeroE6 cells pre-treated with either vehicle control or rISG15 (5µg/mL). D) Infectious viral titer following RSV infection (MOI=0.1, 16h) of VeroE6 cells in the presence of either control IgG or anti-ISG15 antibody (Ab). E) VeroE6 cells pre-treated with either control IgG or anti-ISG15 antibody were infected with mkate2-RSV (MOI=0.1, 48h). WCL collected from the cells were subjected to immunoblotting with RFP (to detect mKate2) and actin antibodies. \**p* < 0.05 & \*\**p* < 0.01 using a Student’s t-test. The immunoblots are representative of data from three independent experiments with similar results.

RSV does not encode a de-ISGylase protein like the PLpro of SARS-CoV-2. Therefore, in contrast to RSV, treatment of Calu-3 and VeroE6 cells with extracellular ISG15 failed to alter SARS-CoV-2 replication (Supp. Fig. 5A, 5B). Since ISGylation confers antiviral activity, we postulate that de-ISGylase activity of PLpro prevents extracellular ISG15 mediated ISGylation-dependent antiviral responses against SARS-CoV-2.

Like IFNs, extracellularly released ISG15 may restrict virus spreading by priming uninfected bystander cells into an antiviral state to ensure reduced viral replication following subsequent infection. Therefore, we investigated whether extracellular ISG15 released from infected cells restricts virus infection and spread. To assess the function of endogenously released extracellular ISG15 during virus infection, we blocked the pro-ISGylation activity of extracellular ISG15 with an anti-ISG15 polyclonal antibody. Addition of the ISG15 antibody, but not the IgG isotype control, reduced induction of intracellular ISG15 protein and ISGylation mediated by extracellular ISG15 (Supp. Fig. 6). Therefore, we used the extracellular ISG15 blocking antibody to examine the antiviral role of ISG15 released during virus infection of IFN-deficient VeroE6 cells. Viral titer analysis revealed approximately 15-fold more infectious virus in ISG15 antibody treated cells compared to control IgG antibody treated cells (Fig. 5D). Concomitantly, we detected higher levels of mKate2 in ISG15 antibody treated cells compared to control IgG treated VeroE6 cells (Fig. 5E). These results demonstrated that extracellular release of ISG15 during virus infection confers IFN-independent antiviral activity.

### Extracellular ISG15 triggers enhancement of intracellular ISG15 protein levels by reducing intracellular ISG15 protein degradation

Extracellular ISG15 robustly enhanced intracellular ISG15 protein levels via an IFN independent mechanism (Fig. 4A, B). ISG15 is an inducible gene that undergoes transcriptional upregulation by IFN following activation of the JAK-STAT pathway^1–9,47,48^. Therefore, we examined the possibility of extracellular ISG15 triggering transcriptional activation of ISG15 via IFN-independent signaling pathway. While IFN-β induced upregulation of ISG15 mRNA in A549 cells (Supp. Fig. 7A), we did not detect upregulation of ISG15 mRNA by extracellular ISG15 in either IFN-competent A549 or IFN-incompetent U5A cell lines (Supp. Fig. 7B, 7C). This result suggested a possible post-transcriptional regulation of intracellular ISG15 protein by extracellular ISG15.

Basal levels of intracellular ISG15 protein are present in non-stimulated cells and thus, in the absence of transcriptional activation of ISG15 gene, we postulated a scenario whereby extracellular ISG15 may reduce the degradation of this pre-existing pool of intracellular ISG15 protein to substantially accumulate intracellular ISG15 protein levels. To examine this scenario, we measured turnover of intracellular ISG15 protein in vehicle versus extracellular ISG15 treated cells in the presence of protein synthesis inhibitor, cycloheximide (CHX). A549 cells were pre-treated with extracellular ISG15 for 2h prior to CHX chase for 0h-6h. While basal intracellular ISG15 levels in vehicle-treated cells rapidly declined, there was distinct stabilization of intracellular ISG15 in extracellular ISG15 treated cells (Fig. 6A). In control vehicle treated cells, half-life of ISG15 was approximately 4h (Fig. 6A, 6B). In contrast, we observed markedly prolonged half-life of ISG15 in extracellular ISG15 treated cells (Fig. 6A, 6B). Only nominal reduction in ISG15 levels was noted in cells treated with extracellular ISG15 and CHX (Fig. 6A, 6B). This phenomenon is specific for extracellular ISG15 since ISG15 protein turnover rate in IFN-β treated cells was approximately 3h (Fig. 6C, 6D), which is similar to turnover of basal ISG15 protein pool in unstimulated control cells (Fig. 6A, 6B). Similar to A549 cells, extracellular ISG15 also markedly stabilized intracellular ISG15 levels in IFN-incompetent U5A cells (Fig. 6E, 6F). In contrast, vehicle treated U5A cells showed rapid turnover of basal ISG15 protein with a half-life of approximately 6h (Fig. 6E, 6F). Profound stabilization of intracellular ISG15 protein by extracellular ISG15 in both IFN-competent (A549) and IFN-incompetent (U5A) cells underscores the intracellular ISG15 protein stabilizing activity of extracellular ISG15 via an IFN-independent mechanism.

**Figure 6.**
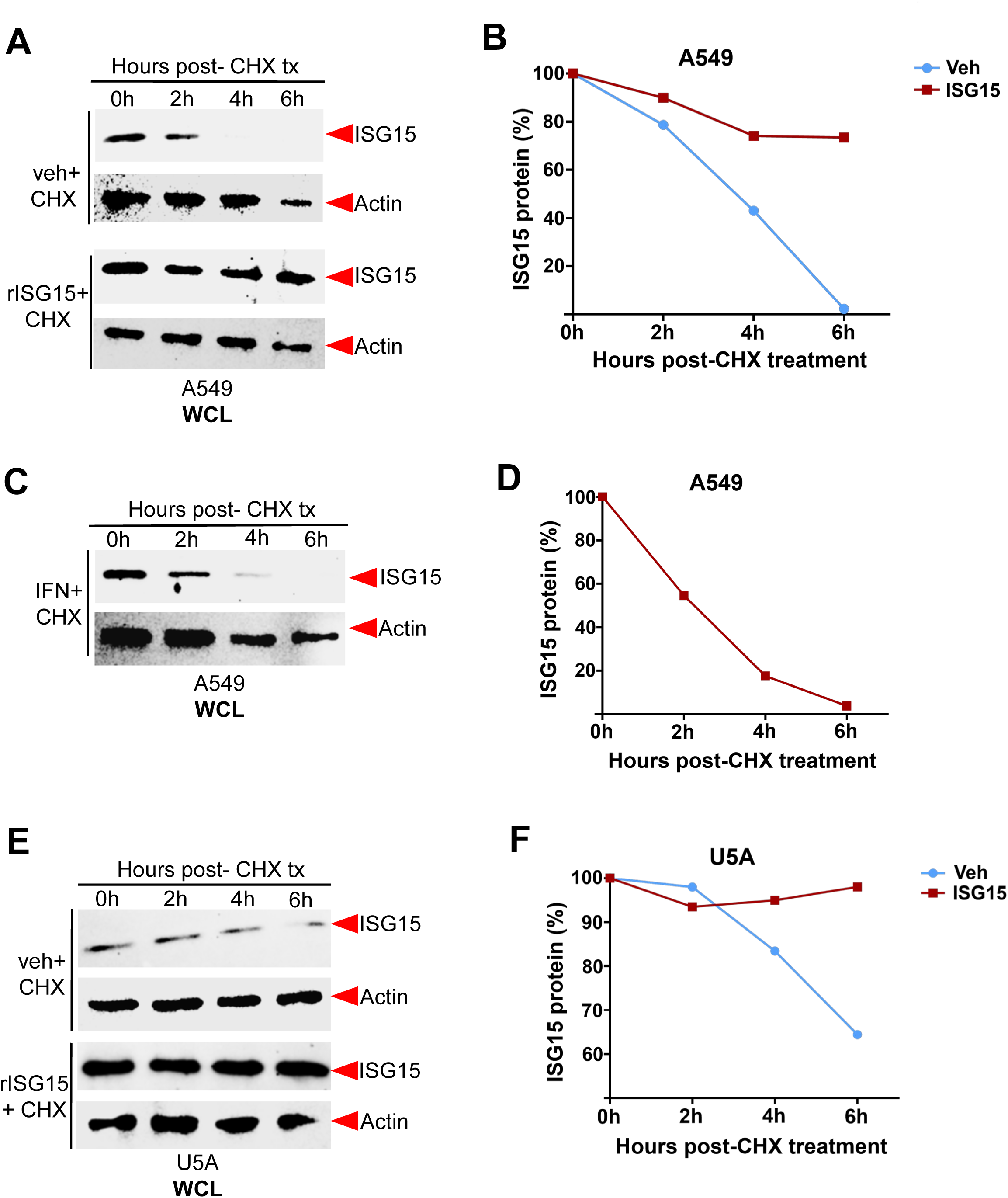
Extracellular ISG15 reduces the turnover of intracellular ISG15 protein by enhancing ISG15 protein stabilization. A) A549 cells were pre-treated with purified recombinant ISG15 protein (rISG15) or vehicle (Veh) control for 2h prior to addition of cycloheximide (CHX). CHX chase (indicated as post-CHX treatment or tx) was performed from 0h-6h and the collected WCL was subjected to immunoblotting with ISG15 and actin antibodies. The vehicle treated blot was over-exposed compared to rISG15 treated blot so that the turnover of basal intracellular ISG15 protein could be clearly analyzed. B) The turnover rate of ISG15 in CHX treated A549 cells was deduced by densitometric analysis of the immunoblot band corresponding to ISG15 from Fig. 6A. The maximum threshold for percent ISG15 protein at 0h was set at 100%. C) A549 cells were pre-treated with interferon-β (IFN) (500 units/ml) prior to addition of CHX. CHX chase was performed from 0h-6h and the collected WCL was subjected to immunoblotting with ISG15 and actin antibodies. D) The turnover rate of ISG15 in CHX treated A549 cells was deduced by densitometric analysis of the immunoblot band corresponding to ISG15 from Fig. 6C. The maximum threshold for percent ISG15 protein at 0h was set at 100%. E) Type I interferon incompetent U5A cells were pre-treated with rISG15 or vehicle (Veh) control for 2h prior to addition of CHX. CHX chase was performed from 0h-6h and the collected WCL was subjected to immunoblotting with ISG15 and actin antibodies. The vehicle treated blot was over-exposed compared to rISG15 treated blot so that the turnover of basal intracellular ISG15 protein could be clearly analyzed. F) The turnover rate of ISG15 in CHX treated U5A cells was deduced by densitometric analysis of the immunoblot band corresponding to ISG15 from Fig. 6E. The maximum threshold for percent ISG15 protein at 0h was set at 100%. The immunoblots are representative of data from three independent experiments with similar results.

Thus, we have identified a yet unknown post-translational regulation of intracellular ISG15 protein via its stabilization from degradation. In the absence of transcriptional upregulation of ISG15 mRNA, we show that intracellular ISG15 levels can be enhanced post-translationally by reducing its degradation. This non-canonical mechanism is utilized by extracellular ISG15 to enhance intracellular ISG15 protein levels.

### Extracellular ISG15 triggers ISGylation via integrin-FAK pathway: interaction of ISG15 with α5β1 integrin

To elucidate the mechanism by which extracellular ISG15 mediates IFN-independent ISGylation, we aimed to identify cell surface molecules that engage with extracellular ISG15. We conducted a preliminary mass spectrometry experiment to identify potential cell surface molecules interacting with extracellular ISG15. For this experiment, we biotinylated purified ISG15 protein (Biotinylated-ISG15). Pre-chilled human lung epithelial A549 cells were incubated with biotinylated-ISG15 at 4°C to prevent internalization of extracellular ISG15. The cell lysate was then precipitated with avidin-agarose, and the bound proteins were subjected to mass spectrometry. Mass spectrometry analysis revealed the interaction of extracellular biotinylated-ISG15 with various cell surface molecules of A549 epithelial cells, including the β1integrin (data not shown). A previous study has shown extracellular ISG15 interacting with integrin molecule LFA-1 in immune cells^28^. Given that LFA-1 is expressed only in immune cells and is not expressed in non-myeloid epithelial cells like A549, we chose the β1 integrin for further analysis since, like LFA-1, it belongs to the integrin family of proteins. Because cell surface integrins are signaling receptors that activate the integrin-focal adhesion kinase (FAK) pathway, we focused our studies on the possibility of extracellular ISG15 interacting with integrin to activate FAK signaling, resulting in IFN-independent ISGylation.

Functional integrin molecules exist as heterodimers of α and β subunits^49–51^. Our mass spectrometry analysis identified β1 integrin as a putative extracellular ISG15 interacting factor and therefore, we investigated whether α5β1 integrin interacts with extracellular ISG15. We confirmed the interaction of extracellular ISG15 with the α5β1 integrin by incubating chilled A549 cells with purified ISG15 protein at 4°C, followed by immunoprecipitation of ISG15 bound proteins with ISG15 antibody and immunoblotting with α5 integrin antibody. Co-immunoprecipitation assay revealed interaction of extracellular ISG15 with α5 integrin (Fig. 7A). To investigate whether ISG15 interacts directly with α5β1 integrin we biotinylated purified α5β1integrin protein. A cell free *in vitro* interaction assay was performed by incubating purified ISG15 protein with biotinylated α5β1integrin (Fig. 7B). We incubated the purified proteins at 4°C, followed by immunoprecipitation with avidin-agarose beads, and immunoblotting with ISG15 antibody. Our cell-free assay demonstrated direct binding of ISG15 protein with α5β1 integrin (Fig. 7B).

**Figure 7.**
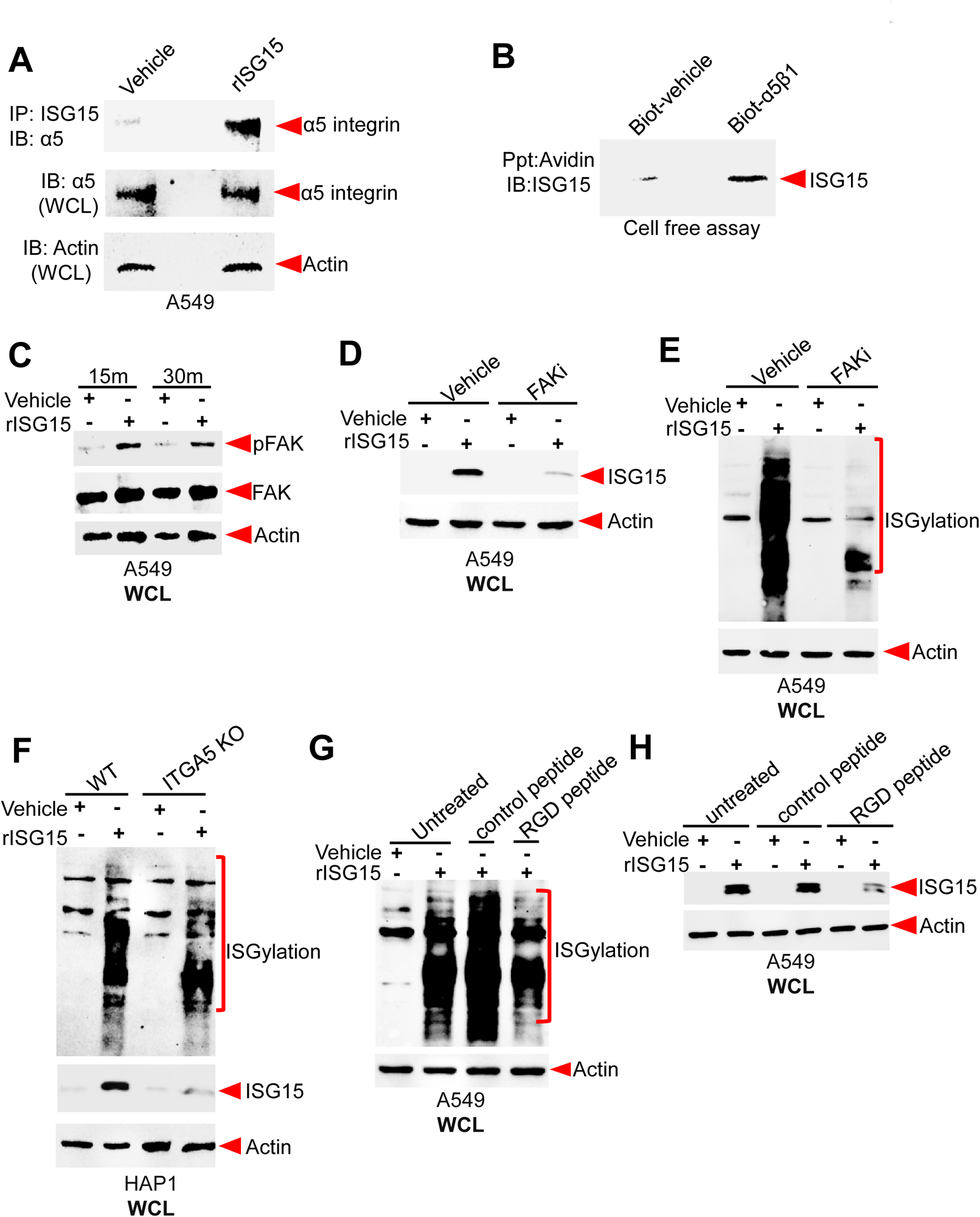
Extracellular ISG15 induces intracellular ISG15 and ISGylation via integrin-FAK pathway: interaction of ISG15 with α5β1 integrin. A) Cell lysates from A549 cells incubated with purified recombinant ISG15 protein (rISG15, 5µg/mL) or vehicle for 4h at 4°C were immuno-precipitated (IP) with ISG15 antibody and subsequently immunoblotted (IB) with α5 integrin antibody. WCL were also blotted with α5 integrin and actin antibodies. B) Biotinylated purified α5β1 integrin protein (5.7 mM) or biotinylated vehicle (control) was incubated with rISG15 protein (0.1 µM) for 16h at 4°C. Following incubation, biotinylated molecules were precipitated (Ppt) with avidin-agarose beads and the avidin-agarose bound complex was immunoblotted (IB) with ISG15 antibody. C) WCL from A549 cells incubated with either rISG15 (1µg/mL) or vehicle for 15 minutes (15m) or 30m were subjected to immunoblotting with phospho-FAK (detects Tyr925 phosphorylation of activated FAK), FAK, and actin antibodies. D) A549 cells were treated with rISG15 (1µg/mL) in the presence of either vehicle control or FAK inhibitor (FAKi, 20 µM). WCL collected from these cells were subjected to immunoblotting with ISG15 antibody to detect unconjugated monomeric form of ISG15 and actin antibody. E) A549 cells were treated with rISG15 (1µg/mL) in the presence of either vehicle control or FAK inhibitor (FAKi, 20 µM). WCL collected from these cells were subjected to immunoblotting with ISG15 antibody to detect ISGylated proteins and actin antibody. F) Wild type (WT) and α5 integrin KO (ITGA5 KO) HAP1 cells were treated with rISG15 (5ug/mL). WCL collected from these cells were subjected to immunoblotting with ISG15 and actin antibodies. G) A549 cells were treated with rISG15 (2µg/mL) in the presence of either control peptide or RGD peptide (200 µM). Cells were also treated with rISG15 in the absence of any peptide (untreated cells). WCL collected from these cells were subjected to immunoblotting with ISG15 antibody to detect unconjugated monomeric form of ISG15 and actin antibody. H) A549 cells were treated with rISG15 (2µg/mL) in the presence of either control peptide or RGD peptide (200 µM). Cells were also treated with rISG15 in the absence of any peptide (untreated cells). WCL collected from these cells were subjected to immunoblotting with ISG15 antibody to detect ISGylated proteins and actin antibody. The immunoblots are representative of data from three independent experiments with similar results.

Activation of cell surface integrins results in transmission of downstream cellular signaling events^49–53^. To elucidate the downstream signaling pathway linked to ISG15-integrin interaction, we investigated the activation status of FAK^54^. FAKs are critical cytosolic adaptors that associate with integrins to mediate signaling following its activation by phosphorylation^54^. Therefore, we examined whether outside-in activation of surface integrins by extracellular ISG15 results in phosphorylation and activation of FAK. Treatment of A549 cells with purified ISG15 protein (rISG15) indeed led to phosphorylation of FAK as detected by immunoblot analyses with phospho-FAK antibody (Fig. 7C). This result suggested the possibility of integrin-FAK dependent activity of extracellular ISG15. The role of the integrin-FAK pathway in extracellular ISG15 mediated activity was assessed by using a FAK inhibitor (FAKi). Profound reduction of intracellular ISG15 protein levels (Fig. 7D) and ISGylation (Fig. 7E) was observed in A549 cells treated with rISG15 protein in the presence of the FAKi. FAKi treatment of IFN-signaling incompetent U5A cells also resulted in reduced intracellular ISG15 protein levels and ISGylation following rISG15 treatment (Supp. Fig. 8A, 8B), indicating that FAK is involved in extracellular ISG15 mediated response in IFN-pathway deficient cells. These results show that the integrin-FAK pathway is a signaling component of IFN-independent ISGylation mediated by extracellular ISG15.

To corroborate the role of integrin signaling during extracellular ISG15 mediated response, we next utilized α5 integrin knockout (ITGA5 KO) human HAP1 cells generated by CRISPR-Cas9 technology^55,56^. Compared to wild-type HAP1 cells, treatment of α5 integrin KO cells with rISG15 led to diminished intracellular ISG15 protein levels and ISGylation (Fig. 7F), confirming the role of integrin during extracellular ISG15 mediated response. We further validated these results using an RGD (Arginine-Glycine-Aspartic acid or Arg-Gly-Asp) peptide to block ligand-integrin interaction. Extracellular matrix (ECM) proteins (e.g., fibronectin, vitronectin, fibrinogen, and von Willebrand factor) utilize the RGD motif to interact with cell surface integrins. Thus, RGD peptides can block the integrin ligand binding site and impair interactions with extracellular ECM ligands and subsequent downstream signaling^57,58^. To determine whether extracellular ISG15 also interacts with integrins similar to RGD-motif containing ECM proteins, we treated A549 cells with rISG15 in the presence of either a control peptide or RGD peptide. Compared to control peptide, RGD peptide treatment resulted in reduced intracellular ISG15 protein levels in extracellular ISG15 treated cells (Fig. 7G). Concomitantly, the RGD peptide also diminished extracellular ISG15 mediated ISGylation (Fig. 7H). The RGD peptide also reduced extracellular ISG15 dependent induction of intracellular ISG15 protein and ISGylation in IFN-incompetent U5A cells, confirming that this mechanism is IFN-independent (Supp. Fig. 8C). This study shows that ISG15 interacts with integrins like other RGD-motif containing integrin ligands. Our *in silico* molecular modeling studies described below provide additional mechanistic insights on ISG15-integrin interaction via RGD-like motifs in ISG15 protein (Fig. 8).

**Figure 8.**
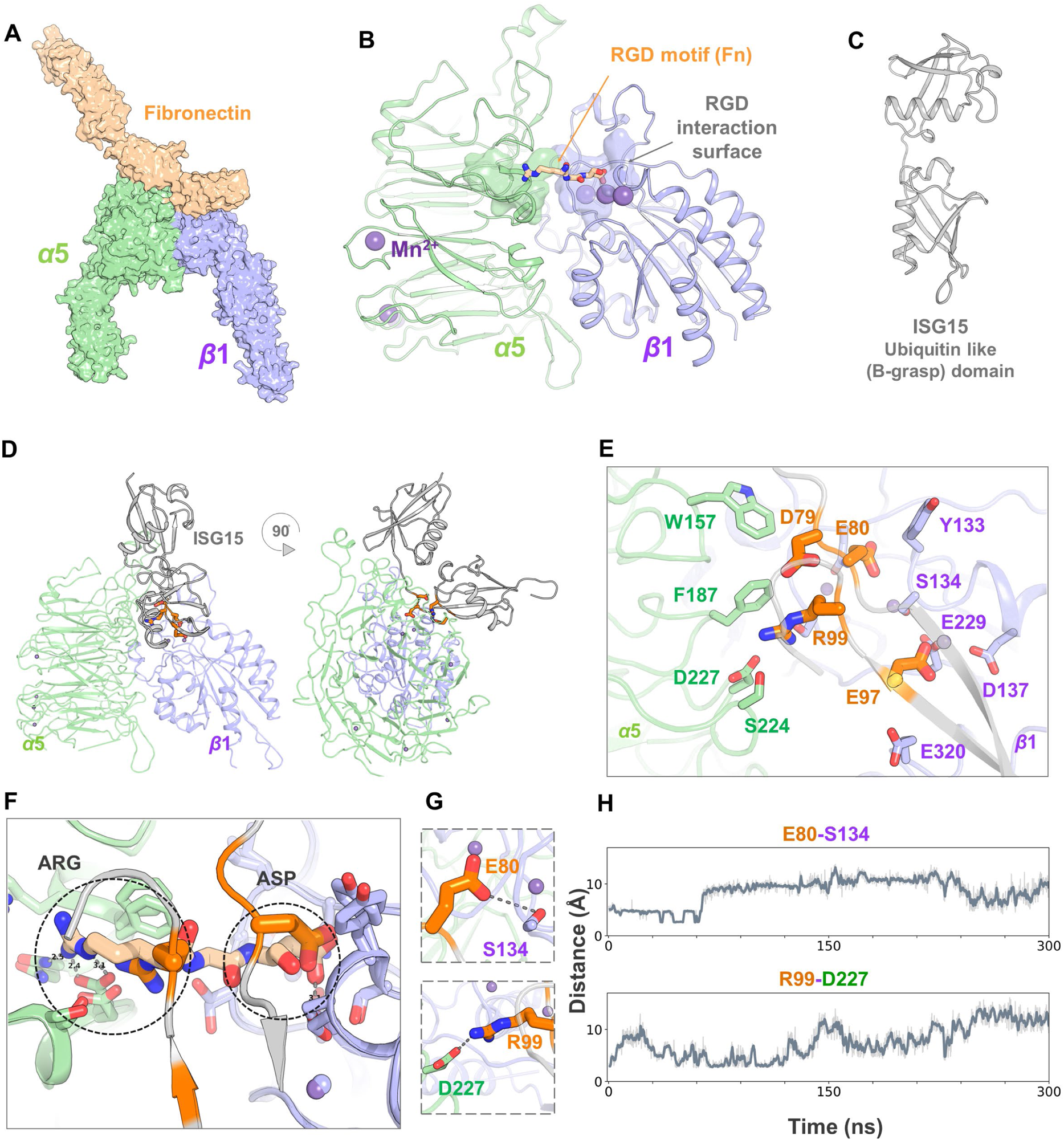
Molecular modeling of extracellular ISG15-α5β1 integrin interaction. A) Crystal structure of α5β1 integrin bound with fibronectin (PDB ID: 7NWL). B) RGD motif binding site at the integrin α5β1 integrin headpiece. RGD residues of fibronectin are shown as licorice, while β-propeller of α5 and βI domain of β1 subunits are shown as cartoon representation. C) Crystal structure of ISG15 (PDB ID: 3RT3). D) Docked orientation of ISG15 with integrin α5β1. E) Residues of ISG15 binding with the RGD motif binding site of α5β1 integrin are shown as orange-colored sticks. F) Superposition of fibronectin RGD residues with the residues of ISG15. Arg99 of ISG15 is positioned similar to the arginine (RGD) of fibronectin and Glu80 occupies the aspartic acid (RGD) binding site. G) Integrin α5β1’s activation-specific interactions with ISG15. H) The two residue-residue interaction distances were monitored throughout the simulation time of 300 ns.

### Molecular modeling of extracellular ISG15-α5**β**1 integrin interaction

Integrin receptors are composed of two types of transmembrane subunits, α and β, forming heterodimeric molecules. Through various combinations of α and β subunits, integrins exhibit multiple subtypes and diverse ligand specificity, enabling them to interact with different ECM molecules^59^. The α5β1 integrin preferentially binds to and is activated by ECM ligands containing the RGD motif, such as fibronectin (Fig. 8A, 8B)^60^.The RGD motif of fibronectin binds within the cleft formed at the interface of the α5 and β1 subunits of integrin. The arginine (Arg) of the RGD motif binds to the α5 subunit, while the aspartic acid (Asp) binds to the β1 subunit, thereby bridging the α-β subunits of integrin (Fig. 8B). Earlier studies on RGD-binding integrins reported that specific structural changes associated with receptor activation occur only in the β subunit of integrins, and that the presence of the aspartic acid within the RGD motif alone is sufficient to activate the receptor^60,61^. Additionally, a recent cryo-EM structure report showed that instead of aspartic acid (Asp), integrin α6β1 recognizes ECM ligand laminin through a glutamic acid (Glu) residue that binds to the β1 subunit of the integrin receptor^62^.

Based on this evidence, we analyzed surface-exposed Arg-Asp and Arg-Glu residue combinations on ISG15 as the potential integrin binding hotspots. ISG15 is comprised of two Ubiquitin-like β-grasp domains (Fig. 8C)^8^. Within this domain, ISG15 has five different sites with exposed Arg-Asp/Glu residue combinations on its surface (Supp. Fig. 9). Each of these five sites of ISG15 was docked individually with the RGD binding site of integrin α5β1 for analysis of activation-specific interactions. Among the docked sites, Site-3 (Supp. Fig. 9) comprised of the residues Arg99 (R99), Glu97 (E97), Asp79 (D79), and Glu80 (E80) exhibited interactions like the known RGD ligands of integrins such as fibronectin (Fig. 8D, 8E). The optimal binding orientation of ISG15 (Site-3) has a HADDOCK docking score of -165.6, which is higher than the scores observed for the other sites of ISG15 (Supp. Table. S1). Surprisingly, we observed Glu80 binding to the aspartic acid (Asp) binding groove of the integrin rather than the typical aspartic acid residues (Fig. 8F). Our analysis suggests that ISG15-α5β1 complex is formed by an RGD-like motif in ISG15, wherein the arginine site of the RGD motif of ISG15 is occupied by Arg99 and the aspartic acid site is occupied by Glu80 (Fig. 8F, 8G). Thus, our analysis has uncovered a new “non-classical” RGD-like motif in ISG15 that can be utilized to achieve similar conformation-driven interaction with integrin similar to ECM ligands harboring “classical” RGD motif.

To further examine the stability of the ISG15-α5β1 interaction, we performed molecular dynamics (MD) simulations of the complex for 300 ns. Overall, the ISG15-integrin α5β1 complex remained stable throughout the simulation time (Fig. 8H).

We also analyzed the stability of the specific interactions that reflect the RGD motif interactions. As it applies to ISG15 – integrin α5β1 complex, Glu80 in the aspartic acid binding groove formed a hydrogen bond interaction with Ser134 of α5β1 (Fig. 8G). The distance between the Glu80 of ISG15 and Ser134 of α5β1 interaction initially starts below 3 Å and remained stable for 80 ns, and it shows a gradual increase in distance over time, stabilizing at around 6 Å (Fig. 8H). Conversely, the interaction between Arg99 of ISG15 and Asp227 of α5β1 shows more variation in distance, oscillating between approximately 3 Å and 10 Å over the simulation period. The Arg99 interaction may be transient and weaker, with the bond forming and breaking throughout the simulation. Upon comparison of the interactions involving Arg99 and Glu80 with the integrin, it appears that ISG15 may interact and activate integrin α5β1 through the stable interaction mediated by Glu80.

Our studies have provided novel insights on innate immune host-defense mechanism by identifying extracellular ISG15 as a new soluble antiviral factor and delineating a type-I IFN independent mechanism for ISG15’s antiviral activity. We show interaction of extracellular ISG15 with cell surface integrins activating integrin-FAK pathway to mediate IFN-independent ISGylation. Additionally, we highlight intracellular ISG15 stabilization as a post-translational mechanism to accumulate ISG15 for ISGylation during IFN-independent signaling. Thus, we put forward a non-canonical mechanism that disengages the widely held dogma about ISGylation being invariably linked with type-I IFN response.

## Discussion

Type-I IFNs are well established antiviral cytokines that confer an antiviral state by activating the JAK-STAT pathway^1–9,29–34,47,48^. One of the mechanisms by which IFNs trigger antiviral response is by promoting ISGylation by inducing expression of IFN stimulated gene 15 (ISG15) protein^3–13,15–26^. ISGylation is an antiviral, post-translational modification wherein intracellular ISG15 is conjugated to target proteins by enzymes related to the ubiquitination process^3–13,15–26^. Although ISGylation is canonically dependent on IFN, our study reports a non-canonical IFN-independent mechanism of ISGylation. We show that extracellular ISG15 activates the integrin-FAK pathway to induce IFN-independent ISGylation. Furthermore, we have identified extracellular ISG15 as a new soluble antiviral factor restricting virus infection via an IFN-independent mechanism. Thus, our study highlights a novel mechanism of IFN-independent ISGylation linked to antiviral activity and mediated by extracellular ISG15 released from virus infected cells.

ISG15 is an inducible gene that is transcriptionally upregulated by IFN-mediated activation of the JAK-STAT pathway^1–9,29–34,47,48^. Our study shows that in the absence of transcriptional activation of the ISG15 gene, extracellular ISG15 can substantially enhance the stability of intracellular ISG15 protein to ensure abundant ISG15 is available for ISGylation. This highlights a new IFN-independent non-canonical post-translational mechanism regulating intracellular ISG15 levels.

Antiviral activity of ISGylation restricts infectivity of a wide-range of RNA and DNA viruses including Ebola virus, IAV, rabies virus, Zika virus, dengue virus, WNV, LCMV, HIV-1, HBV, HCV, CVB3, Sindbis virus, KSHV, HSV-1, CMV^15–26^. Therefore, we envision that apart from RSV and SARS-CoV-2, other viruses may also trigger release of ISG15 during infection to launch host-defense apparatus to counteract infectivity via non-canonical ISGylation by extracellular ISG15. Moreover, ISG15 and ISGylation regulate two key innate antiviral mechanisms including STING-cGAS and MDA5 antiviral pathways^35,63–65^. Thus, our study has wide ramification in understanding innate antiviral mechanisms during infection with a wide variety of viruses.

Apart from viral innate immunity, ISGylation plays an role during various non-viral diseases such as bacterial diseases (Listeria monocytogenes, tuberculosis), non-infectious diseases such as cancer (breast, lung, colon, pancreatic, prostate cancers), and neurodegenerative disorders (Amyotrophic lateral sclerosis or ALS, Parkinson’s disease, multiple sclerosis or MS, Ataxia telangiectasia, brain injury)^14^. We have identified a non-canonical mechanism of ISGylation by extracellular ISG15, which may have a role during non-viral diseases. Particularly, ISG15 is released from tumors and extracellular ISG15 and ISGylation promotes tumorigenesis^14,66–69^. Therefore, blocking non-canonical ISGylation by extracellular ISG15 may serve as a therapeutic target to combat diseases such as cancer. Thus, our study has wider implication in understanding pathogenesis of not only viral diseases, but non-infectious diseases like cancer and neurological disorders.

Previous studies have reported release of ISG15 from macrophages infected with SARS-CoV-2 and influenza A virus (IAV)^36^. Bronchial epithelial cells infected with rhinovirus also released ISG15 into the extracellular milieu^37^. Extracellular ISG15 did not reduce rhinovirus infection, but instead downregulated the immune response by inhibiting the release of the chemokine CXCL10 from infected cells^37^. Circulating ISG15 has been detected in serum of patients infected with SARS-CoV-2^35^, and levels of free ISG15 in the saliva and nasal secretion of rhinovirus patients correlated with viral load^37^. Although extracellular ISG15 has been detected in clinical samples of virus-infected patients, the functional role of extracellular ISG15 during virus infection has not been reported, and the role of viral encoded factor(s) in ISG15 release is unknown. Previous reports have identified the SARS-CoV-2 S protein in upregulating the expression of *isg15*^39^. Our study has identified the S protein of SARS-CoV-2 and other coronaviruses as a viral factor involved in induction of ISGylation and in ISG15 release.

SARS-CoV-2 encodes a de-ISGylase (PLpro) that removes conjugated ISG15 from its targets, negating the antiviral effects of ISGylation^35,70^. Therefore, ISGylation has yet to be detected in SARS-CoV-2 infected cells. Although ISGylation was not observed during early-mid infection, interestingly, we detected ISGylation and release of ISG15 from SARS-CoV-2 infected human lung epithelial cells during late infection (Fig. 1A, 1B). We further show that SARS-CoV-2 PLpro expression in lung epithelial cells during early-mid infection correlated with lack of ISGylation (Fig. 1A). However, PLpro declined during late infection, when ISGylation and ISG15 release were evident (Fig. 1A, 1B). Due to the de-ISGylase activity of PLpro, extracellular ISG15 mediated ISGylation did not reduce SARS-CoV-2 replication (Supp. Fig. 5). However, based on the activity of extracellular ISG15 on Natural Killer cells (NK cells)^28^, it is possible that ISG15 released from infected epithelial cells during SARS-CoV-2 infection acts on immune cells like macrophages to shape inflammatory and adaptive responses. Indeed, it has been speculated that ISG15 release during SARS-CoV-2 contributes to hyperinflammation via uncharacterized mechanisms^71^. On the other hand, RSV does not encode a de-ISGylase, and ISGylation is effective at limiting RSV infection^42^. We show a direct effect of extracellular ISG15 in promoting the antiviral response in epithelial cells by restricting RSV infection. RSV infection led to ISG15 release (Fig. 2B) presumably by expression of RSV F protein since ectopic expression of F triggered ISGylation and ISG15 release (Fig. 2E). Extracellular ISG15 released during RSV infection induced ISGylation in an IFN-independent manner to promote an antiviral response, which reduced RSV infection (Fig. 4C, 4D, 4E, 5). We postulate that antiviral activity of circulating ISG15 in the extracellular milieu may dictate disease outcome and secondary complications during virus infection. Therefore, viruses like SARS-CoV-2 have evolved to encode viral proteins like PL-pro to antagonize the antiviral effect of extracellular ISG15 by blocking ISGylation. In contrast, ISGylation mediated antiviral activity by extracellular ISG15 cannot be counteracted by viruses like RSV that do not possess any viral tools to antagonize ISGylation. Furthermore, we show a key role of viral envelope proteins like S and F proteins in promoting ISGylation and ISG15 release.

ISG15 release has also been documented in non-viral settings, including treatment of myeloid cells (NK cells, PBMCs, T-cells) with various pathogen associated molecular patterns (PAMPs) and over-expression of ISG15 in HEK293 cells^27^. Extracellular ISG15 released from HEK293 cells interacted with LFA-1 integrins on NK cells and induced the release of IFNγ, but only when combined with IL-12^28^. Contrasting these studies, we show that extracellular ISG15 alone elicited induction of intracellular ISG15 protein, ISGylation, and antiviral response in epithelial cells. Our data and that in myeloid cells highlights cell-dependent immunomodulatory activities by extracellular ISG15 dictated by its interaction with two unrelated cell surface integrins: LFA-1 in myeloid cells^28^ and α5β1 in non-myeloid epithelial cells (our study).

In immune myeloid cells, extracellular ISG15 interacts with the integrin LFA-1 and signals through intracellular Src proteins, leading to the release of IFNγ^28^ in IL-12 primed NK cells^72^. However, we have identified pro-ISGylation and antiviral activity of extracellular ISG15 in non-myeloid epithelial cells. In that regard, LFA-1 is only expressed in myeloid cells, and thus the receptor involved in transducing the activity of extracellular ISG15 in non-myeloid cells was unidentified. Although epithelial cells do not express LFA-1, other integrins are expressed at their surface. Among the epithelial surface integrins, α5β1 integrins are highly abundant in lung tissue. α5β1 integrins are involved in tissue remodeling and respiratory diseases^73^. Recently, a role of α5β1 integrins in regulating pro-inflammatory response by respiratory viruses and TNF-α via oxysterols was reported^56,74^. α5β1 integrins participate in cell survival and adhesion by binding to RGD (Arginine, Glycine, Aspartic Acid) motif containing proteins like fibronectin within the extracellular matrix (ECM). To do this, α5β1 integrin adopts a half-bent conformation. This leaves its RGD binding domain partially exposed to interact with RGD containing ECM proteins^60^. RGD peptide blocked ISGylation activity of extracellular ISG15 (Fig. 7G, 7H) (Supp. Fig. 8C), and accordingly, we show that like other integrin ligands, ISG15 has an RGD-like domain (Fig. 8) that facilitates interaction with α5β1 to trigger outside-in signaling^75^. Our *in silico* molecular docking studies (Fig. 8) show that ISG15 can interact with the RGD binding region of α5β1 by utilizing Arginine and Glutamic acid amino acids to mimic the Arginine and Aspartic acid residues of the RGD motif. Indeed, extracellular ISG15 interacted with α5β1 integrin in a cell free interaction assay and with the epithelial cell surface α5β1 integrin (Fig. 7A, 7B). This type of interaction is similar to that between extracellular ISG15 and LFA-1^28^. ISG15/LFA-1 interaction culminated in IFNγ production (combined with IL-12) from myeloid cells, while our study demonstrated ISG15/α5β1 integrin interaction leading to intracellular ISGylation independent of the canonical IFN-dependent mechanism.

FAKs are the main mediators of integrin signaling^54^. Ligands bind to integrins which rearrange into focal adhesions, leading to activation of submembrane FAK molecules by its autophosphorylation^54,75^. For example, fibronectin, which has an RGD domain, can bind to integrins and activate STAT1 in a mechanism dependent on activated FAK^76^. FAK also forms complexes with downstream adapters, like Src, to continue a phosphorylation cascade to activate other proteins ^77^. We show FAK activation by extracellular ISG15 (Fig. 7C). In addition, when FAKs are chemically blocked prior to extracellular ISG15 treatment, ISGylation is impaired (Fig. 7D, 7E) (Supp. Fig. 8A, 8B), suggesting that the integrin-FAK cascade is involved in extracellular ISG15 mediated ISGylation. FAKs also play a role during virus infection to activate proteins that are associated with different stages of the innate immune response. MAVS-mediated antiviral signaling was shown to be facilitated by FAK localization to the mitochondrial membrane during Sendai Virus (SeV) infection^78^. Phosphatidylinositol 3-kinase (P13K) signaling during influenza A virus (IAV) infection was also found to be regulated by FAK phosphorylation^79^. Thus, FAK participation in ISG15-integrin signaling in promoting an antiviral response is to be considered in future studies.

Several studies have reported IFN-independent transcriptional induction of ISGs via various transcription factors (IRF3, IRF7) and intracellular PAMPS such as dsDNA (human cytomegalovirus or HCMV) and bacterial DNA (*Listeria* and *Mycobacterium*)^19,80–83^. It is unlikely that extracellular ISG15 is utilizing transcription factors and/or PAMPs, since it triggered intracellular ISG15 protein accumulation for ISGylation at a post-translational stage by virtue of reducing intracellular ISG15 protein degradation. Several mechanistic possibilities exist that may confer such stabilizing activity of extracellular ISG15 via integrin-FAK pathway. Numerous studies have demonstrated FAK signaling modulating protein stability^77,84–86^. Protein stability regulation by FAK signaling occurs via subcellular spatial re-localization and/or activation of various scaffolding proteins that form complexes with target proteins to either accelerate or decelerate their degradation^77,84–86^. Apart from FAK, cell surface Frizzled-LRP5/6 receptor signaling activated by extracellular Wnt (wingless) ligands stabilizes cytoplasmic β-catenin protein by preventing its degradation^87–89^. Accumulated β-catenin protein is translocated to the nucleus for transactivation of β-catenin/Wnt dependent genes. Similar to FAK signaling, β-catenin protein stability is also modulated by scaffolding protein complex interacting with cytoplasmic β-catenin protein^87–89^. We envision that a similar apparatus may play a role in intracellular ISG15 protein stabilization following activation of ISG15-integrin-FAK pathway. In the future, we will attempt to identify and characterize the scaffolding proteins that may complex with intracellular ISG15 protein to dictate its degradation rate following activation of integrin-FAK signaling by extracellular ISG15.

Three enzymes (UBE1L, UBCH8 and HERC5) that are required for ISGylation are IFN inducible genes. Therefore, several possibilities may trigger ISGylation by these enzymes during IFN-independent ISGylation by extracellular ISG15. First, existing basal levels of enzymes present in unstimulated cells may be adequate to trigger ISGylation since enzymatic reactions do not require high levels of enzyme proteins. Secondly, similar to intracellular ISG15 protein, extracellular ISG15 may also enhance the stability of these enzymes to ensure ISGylation. Thirdly, a scenario may exist whereby once an adequate intracellular ISG15 protein level threshold is achieved, cells can “sense” this occurrence by a yet to be defined IFN-independent mechanism to transcriptionally induce enzymes required for ISGylation. In the future, we will perform detailed studies to identify the mechanism regulating the dynamics of ISGylation-related enzymes during extracellular ISG15 mediated IFN-independent response.

Extracellular ISG15 may not only bind to α5β1, but a diverse set of integrins on the surface of multiple cell types that in turn may govern additional facets of antiviral activity. Viruses such as Ebola virus^90^, Epstein Barr virus^91^, and foot and mouth disease virus^92,93^ can use α5β1 and other integrins for cellular entry. Extracellular ISG15 may block virus entry by occupying or altering the conformation of integrins to prevent virus from interacting efficiently with integrin receptors for cellular entry. This is an intriguing research avenue for the development of ISG15 derived competitive virus inhibitors and therapeutics or to prevent off-site effects from extracellular ISG15 at distant sites. In summary (Fig. 9), our studies identified ISG15 as a new extracellular antiviral factor that triggers non-canonical ISGylation in an IFN-independent mechanism. Here, extracellular ISG15 interacts with cell surface integrins to trigger FAK-dependent ISGylation by stabilizing intracellular ISG15 protein. Since ISGylation occurs widely during virus infection, this mechanism has broad implications for understanding virus-host interactions and development of host-directed therapeutics against multiple viruses. Furthermore, the non-canonical ISGylation mechanism highlighted in our study may also regulate non-viral ISGylation-associated diseases such as cancer and neurodegenerative disorders.

**Figure 9.**
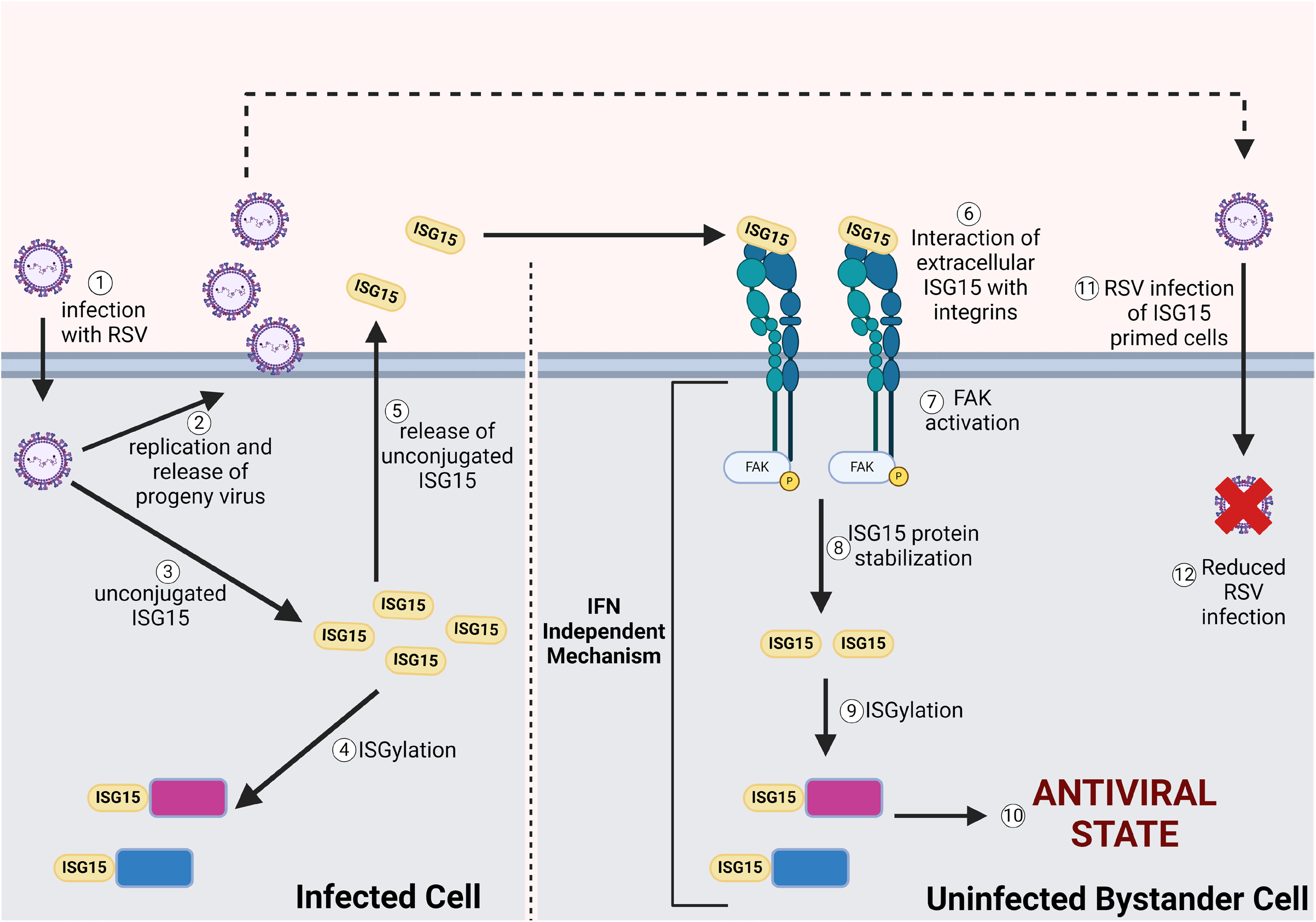
Schematic model showing antiviral pro-ISGylation activity of extracellular ISG15 via a non-canonical type-I interferon (IFN) independent mechanism. Virus infection (1) will trigger induction of intracellular unconjugated ISG15 protein (3), which will be utilized for ISGylation (4). A portion of unconjugated ISG15 will be released from infected cells (5). Extracellular ISG15 will interact with the cell surface integrin molecules (6) of uninfected bystander cells (paracrine action) to trigger FAK activation (7). This will result in post-translational stabilization of intracellular ISG15 protein (8) for ISGylation (9) via an IFN independent mechanism. Since ISGylation possesses antiviral activity, it will confer an antiviral state in the uninfected bystander cells (10) to ensure reduced viral infectivity during subsequent infection of these cells (11 & 12) by progeny infectious virus produced from infected cells (2). Apart from paracrine action, extracellular ISG15 can also act on infected cells (autocrine action) for optimal antiviral response.

## Authorship

L.G.M, K.C, C.M, I.M, S.P, L.M, P.D and T.A.M. performed the experiments. L.G.M, K.C, C.M, A.L.H, P.D, M.N.T, S.N, T.A.M, S.B contributed to the experimental design, data analyses and interpretation, L.G.M, K.C, C.M, P.D prepared the figures and tables, manuscript was written by L.G.M, K.C, C.M, S.N, T.A.M, S.B.

## Acknowledgements & Funding

This research was supported by funding from the Washington Research Foundation (SB), National Institutes of Health (NIH) R01AI083387 (SB), and R01GM137022 (SN). LGM was supported by NIH NIGMS Predoctoral Biotechnology Training Grant 5T32GM008336. Biorender was used to create the model and timelines for this project. We would also like to thank the Franceschi Microscopy and Imaging Center of Washington State University for use of their facilities and staff assistance.

## Materials and Methods

### Cell Culture

A549, Calu-3, and VeroE6 cells were maintained in DMEM (GIBCO #11995065) supplemented with 10% FBS, 100 IU/ml penicillin, and 100 g/ml streptomycin. CV-1, U5A and 2fTGH cells were maintained in DMEM supplemented with 5% FBS, 100 IU/ml penicillin, and 100 g/ml streptomycin. WT and ITGA5 KO HAP1 cells were maintained in IMEM supplemented with 10% FBS and 100 IU/ml penicillin, and 100 g/ml streptomycin.

### Cell Transfection

pcDNA3.1 plasmids encoding the C9-tagged SARS-CoV-2 Spike (S) protein (pcDNA3.1-SARS2-S) (Addgene plasmid # 145032), SARS-CoV S protein, MERS S protein, HCoV-229E S protein were used in these experiments; all were provided by Dr. T. Gallagher (Loyola University, Chicago, IL, USA). pcDNA3.1 plasmid encoding the RSV Fusion (F) protein was generated in the laboratory. Cells were transfected with either coronavirus S proteins, RSV F protein, or empty vector (pcDNA3.1) using Lipofectamine 2000 (Invitrogen) and maintained in antibiotic free DMEM supplemented with 10% FBS. After 24h, the media was changed and the WCL or supernatant was collected at the indicated timepoints. Similar transfection methodology was utilized to express RSV Fusion (F) protein in A549 cells.

### Recombinant S protein, IFN-β, ISG15, FAK inhibitor (FAKi), cycloheximide (CHX), and RGD Blocking Peptide Treatment

Cells were treated with 500 U/ml of IFN-β (Sino Biological, catalog no. 10704-HNAS) or vehicle prepared in antibiotic free DMEM supplemented with 10% FBS. A549, VeroE6, U5A, 2fTGH, HAP1 (WT and ITGA5 KO) cells were also treated with 5µg/mL of recombinant purified ISG15 protein (rISG15) for indicated times before collection of whole cell lysates. The rISG15 protein was purified as described previously^4^ and we also used rISG15 purchased from Sino Biological (catalog no.12729-HNAE1). For CHX experiments, cells were treated with ISG15 (5ug/mL) or IFN (500U/mL) for 2h and then treated with CHX (Sigma-Aldrich catalog no. C7698-1G) (75ug/mL) for the indicated timepoints. For FAK inhibitor experiments, A549 and U5A cells were treated with the FAK inhibitor (PF-431396 hydrate; Sigma-Aldrich, catalog no. PZ0185) at a concentration of 20µM in 10% or 5% antibiotic-free media, respectively. After 2h, 1µg/mL of rISG15 was added to the media containing FAKi. Alternatively, A549 cells were pretreated for 2h using 200µM of RGD blocking peptide (AnaSpec Inc., catalog no. AS-62049) or control peptide before adding rISG15 (2µg/ml) to the media. In some experiments, A549 cells were treated with 8nM of full-length SARS-CoV-2 S protein (BEI catalog no. NR-52397) or the S1 subunit (Sino Biological, catalog no. 40591-V08HB) for 16h before collection of whole cell lysate.

### Immunoblot

Whole cell lysates (WCL) were collected using 1x Laemmli sample buffer containing beta-mercaptoethanol. Lysates were subject to gel electrophoresis in an acrylamide gel and then transferred to Amersham Protran 0.2-um nitrocellulose blotting membranes. Membranes were blocked in 5% milk with PBS-T for 5h at room temperature and blotted with primary antibodies against ISG15 (affinity purified polyclonal ISG15 antibody was generated as described previously^4^), Rhodopsin C9 (Invitrogen, catalog no. MA1-722), S1 (Sino Biological catalog no. 50150-R007), RFP (Thermo Fisher Scientific, catalog no. MA5-15257), pFAK(Y397) (Cell signaling, catalog no. 8556P), FAK (Cell signaling catalog no. 13009S), α5 integrin (Proteintech Catalog no.10569-1-AP), RSV F (RSV F 101F antibody was acquired from Dr. Jason McLellan, University of Texas, Austin, TX, USA), SARS-CoV-2 PLpro (GeneTex catalog no.44328), and actin (Bethyl Laboratories, Inc., catalog no. A300-491A) overnight at 4°C. Secondary antibodies conjugated to horseradish peroxidase were added for 1.5h at room temperature. Protein bands were developed using Western Lightning Plus-ECL (Perkin Elmer catalog no. 104001EA). In some experiments, protein bans were quantified using ChemiDoc XRS software Image Lab 5.1 (Bio-Rad).

### RNA Extraction and RT-qPCR

RNA was extracted using Trizol reagent (Life Technologies, catalog no. 15596026) following the manufacturer’s instructions. For analysis of SARS-CoV-2 nucleocapsid (N) gene levels, RNA was treated with ezDNase and cDNA was synthesized using SuperScript IV VILO Master Mix (Thermo Fisher, catalog no. 11766050). qPCR was performed using PowerUp SYBR Green Master Mix following manufacturer’s instructions (Thermo Fisher; catalog no. A25741). For analysis of ISG15 transcript levels, RNA was treated with DNase (Thermo Scientific, catalog no. ENO521) and cDNA was synthesized using Multiscribe Reverse Transcriptase (Thermo Fisher, catalog no. 4319983). qPCR assay was performed using PerfeCTa SYBR Green FastMix (Quantabio; Catalogue no. 95073-012) using the manufacturer’s instructions. qPCR was performed in a StepOnePlus Real-Time PCR System (Applied Biosystems). Abundance of SARS-CoV-2 N mRNA was normalized to that of GAPDH and ISG15 was normalized to GAPDH and vehicle control by DDCt method. The following primers were used for RT-qPCR:

SARS-CoV-2 N Fw: 5′-CAATGCTGCAATCGTGCTAC-3′

Rv: 5′-GTTGCGACTACGTGATGAGG-3′

Human ISG15 Fw: 5’-CGCAGATCACCCAGAAGATCG-3’

Rv: 5’-TTCGTCGCATTTGTCCACCA-3’ GAPDH

Fw: 5’-ACAACTTTGGTATCGTGGAAGG-3’

Rv: 5’-GCCATCACGCCACAGTTTC-3’

### TCA Precipitation of Proteins from Medium Supernatant

A549 cells were transfected with either S protein, F protein or pcDNA (control) for 24h. Media was changed to remove plasmids and washed cells were cultured for an additional 16h or 24h in plasmid free media. Plasmid-free cell culture medium supernatants were collected and clarified to remove cell debris. Total proteins in medium supernatants were precipitated with Trichloroacetic acid (TCA, 20% v/v final concentration) overnight at 4°C. Protein pellets were washed three times with 200ul of ice-cold acetone. Pellets were resuspended in 2X Laemmli buffer with beta-mercaptoethanol and neutralized with 1M Tris-HCL at pH 8.0 before immunoblotting with indicated antibodies. Similar TCA precipitation and immunoblotting was performed with medium supernatants collected from SARS-CoV-2 and RSV infected A549, Calu-3, and VeroE6 cells.

### Viruses

SARS-CoV-2 isolate USA-WA1/2020 (BEI resources catalog no. NR-52281) was propagated in VeroE6 cells. Human respiratory syncytial virus A2 strain was purified using previous methods^43,56,94,95^. Recombinant RSV expressing mkate2 (RSV-mkate2) was propagated as described previously^41,96^.

### Infection Experiments

Work with infectious SARS-CoV-2 was performed in biosafety cabinets within a biosafety containment level 3 facility. Personnel wore powered air purifying respirators during all procedures (MAXAIR Systems, Irvine, CA). Work with RSV was performed in biosafety cabinets within a biosafety containment level 2 facility. As indicated, A549, VeroE6 and Calu-3 cells were infected with either SARS-CoV-2 (MOI=0.1) or RSV (RSV-mkate2 or RSV at MOI=0.1). After indicated times, medium supernatants were collected for TCA precipitation followed by immunoblotting. Whole cell lysates were collected in Laemmli buffer for immunoblotting. Calu-3 and VeroE6 cells were also pretreated with 5ug/mL of rISG15 for 8h and then infected with SARS-CoV-2 (MOI=0.1) for RNA collection using Trizol. A549 and VeroE6 cells were infected with RSV-mkate2 (MOI=0.1) for 48h. Whole cell lysate was collected in Laemmli buffer for immunoblotting and medium supernatant was collected for TCA precipitation assays. In some experiments, A549 and VeroE6 cells were pretreated with 5µg/mL of rISG15 for 8h, and then infected with either RSV-mkate2 or wt-RSV for 48h. Whole cell lysate of RSV-mkate2 infected samples were collected in Laemmli buffer and subject to immunoblotting. Supernatants of samples infected with wt-RSV were collected for plaque assay to analyze infectious virus titer. For plaque assays, CV-1 cells were infected with ten-fold dilutions of the supernatants. After 1.5h adsorption, methyl cellulose was added to the wells, and incubated for 48h-72h. The methyl cellulose was removed, and the plates were dyed with crystal violet in 70% ethanol. After 45 minutes, the plates were washed with ddH2O. Vero cells were also infected with RSV-mkate2 or wt-RSV for 48h in the presence of either control rabbit IgG (Cell signaling, catalog no. 2729) or affinity purified ISG15 polyclonal antibody^4^. Lysate was subject to immunoblotting, and supernatant was used to perform TCID50 assays. Infectious virus titer was calculated using the Reed and Meunch method.

### LDH Assays

Medium supernatants from mock and virus (RSV or SARS-CoV-2) infected cells was collected at indicated timepoints. LDH assays were performed according to manufacturer’s instructions (Abcam, catalog no. AB65393). To calculate cell death the percentage, we used the calculation: ((test sample-low control)/(high control-low control)) × 100.

### Phase contrast microscopic image

VeroE6 cells seeded on chamber slides were pretreated with 5µg/mL of ISG15 for 8h and then infected with RSV (MOI=1) for 16h. Live cells were imaged using a Leica microsystems microscope (Model DM18).

### Confocal microscopy

VeroE6 cells were infected with RSV-mkate2 (MOI=0.1) for 48h. Infected cultures were washed and fixed in 4% paraformaldehyde, blocked in 5% BSA in PBS-T and incubated overnight with primary antibody against ISG15. Slides were incubated with CoraLite488-conjugated secondary antibody (Proteintech, catalog no. SA00013-2) and mounted with DAPI ProLong Gold antifade (Invitrogen, catalog no. P36930). The fluorescent images were analyzed by using a Leica TCS SP8 X confocal microscope.

### Interaction Assays

For cell-free assays, purified α5β1 integrin protein (Yo proteins, catalog no. 638) was biotinylated by using The EZ-link® TFPA-PEG3-Biotin (Thermo Fisher Scientific cat no 21303) as per manufacturer’s instruction. Biotinylated α5β1integrin (5.7 mM) was incubated with NeutrAvidin Agarose Resin beads (Thermo Scientific, catalog no. 29200) for 16h at 4°C. Purified ISG15 protein (rISG15) was then added to the avidin beads (bound to biotinylated integrin) and incubation was continued for additional 16h at 4°C. 2X Laemmli sample buffer was added to the beads for further immunoblotting analysis.

For experiments with cells, A549 cells were chilled at 4°C for 15 minutes prior to addition of rISG15 (5µg/mL) to chilled A549 cells. After 4h incubation at 4°C, the lysate was subjected to co-immunoprecipitation analysis with α5 integrin and ISG15 antibodies.

### Docking of ISG15 and Integrin α5β1

For modeling the ISG15-integrin α5β1 complex, protein-protein docking was carried out. The crystal structures of ISG15 (PDB ID: 3RT3) and integrin α5β1 (PDB ID: 7NWL) were downloaded from the Protein Data Bank^97^. Missing side-chain atoms and residues were added, and the structure was optimized using the structure preparation module of MOE^98^. For the docking of ISG15 with integrin α5β1, the residues comprising the RGD motif binding site (Tyr186, Phe187, Gln189, Gln221, Ser224, and Asp227 of the α5-β propeller domain, and Ser132, Tyr133, Ser134, Gly223, Asn224, Leu225, Asn226, Ser227, and Glu229 of the β1-βI domain) of α5β1 were selected as the potential interaction site. For ISG15, the exposed arginine and aspartic acid residues (similar to the RGD motif) were used as the possible interaction site. ISG15 has five surface-exposed combinations of arginine with aspartic acid residues. Each site of ISG15 was docked to the α5β1 integrin’s RGD binding site. Protein-protein docking simulations were performed using the HADDOCK v2.4 server^99,100^. For each protein-protein docking run, at most 20,000 possible binding orientations were generated, among which 2000 poses were considered for post-docking minimization. Finally, 1,000 poses were filtered and used for scoring and clustering. Among the resulting clusters with multiple docking poses, a cluster exhibiting interactions resembling those of the RGD motif of integrins was selected for subsequent MD simulation.

### MD Simulation of ISG15 - α5β1 complex

The stability of the ISG15-α5β1 complex was further examined using molecular dynamics simulations using AMBER v20^101^. The input files for the MD simulation were generated using the Input Generator-Solution Builder module of CHARMM-GUI with AMBERff14SB forcefield^102^. The system was solvated using TIP3P water molecules in a cubic box such that the distance between any atom of the protein complex and the box edge was at least 10 Å^103^. Subsequently, the system was neutralized (net charge = 0), and the salt concentration was brought to 0.15 M by adding Na+ and Cl− ions. The simulations were performed under periodic boundary conditions and with the Particle Mesh Ewald (PME) method for calculating the long-range electrostatic interactions^104,105^. The van der Waals interactions were smoothly switched off at 12 A°. Further, the solvated system was minimized (10,000 steps) to remove any steric clashes in the system. Following the minimization step, equilibration and production runs were performed with an integration time step of 2 fs, and all the bond lengths involving hydrogen atoms were fixed using the SHAKE algorithm^106^. The system was equilibrated for 5 ns using an NPT ensemble at 1 atm pressure and 310 K temperature with constraints, the production simulations were carried out for 300 ns without any constraints, and the trajectory was saved for every 10 picoseconds.

### Statistical analysis

Data were analyzed using Graphpad Prism software (6.0) and significance test was carried out using Student’s t-test.

