## Supplementary data for "Extracellular ISG15 triggers ISGylation via a type-I interferon independent non-canonical mechanism to regulate host response during virus infection"

### Supplementary Fig. 1

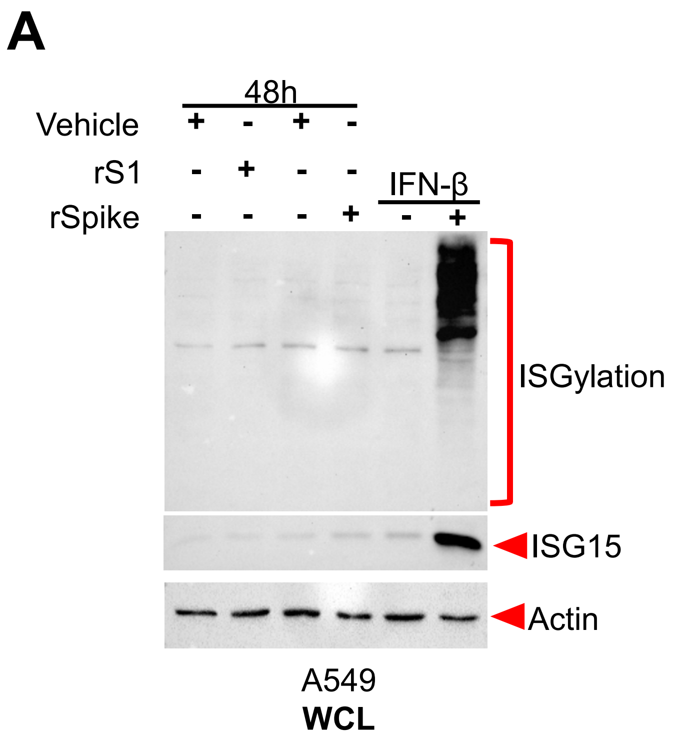


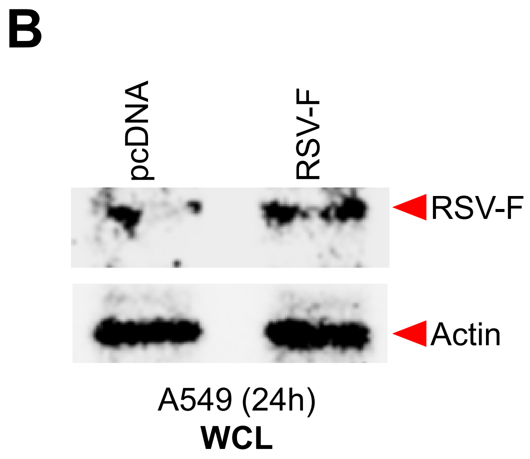

### Supplementary Fig. 2

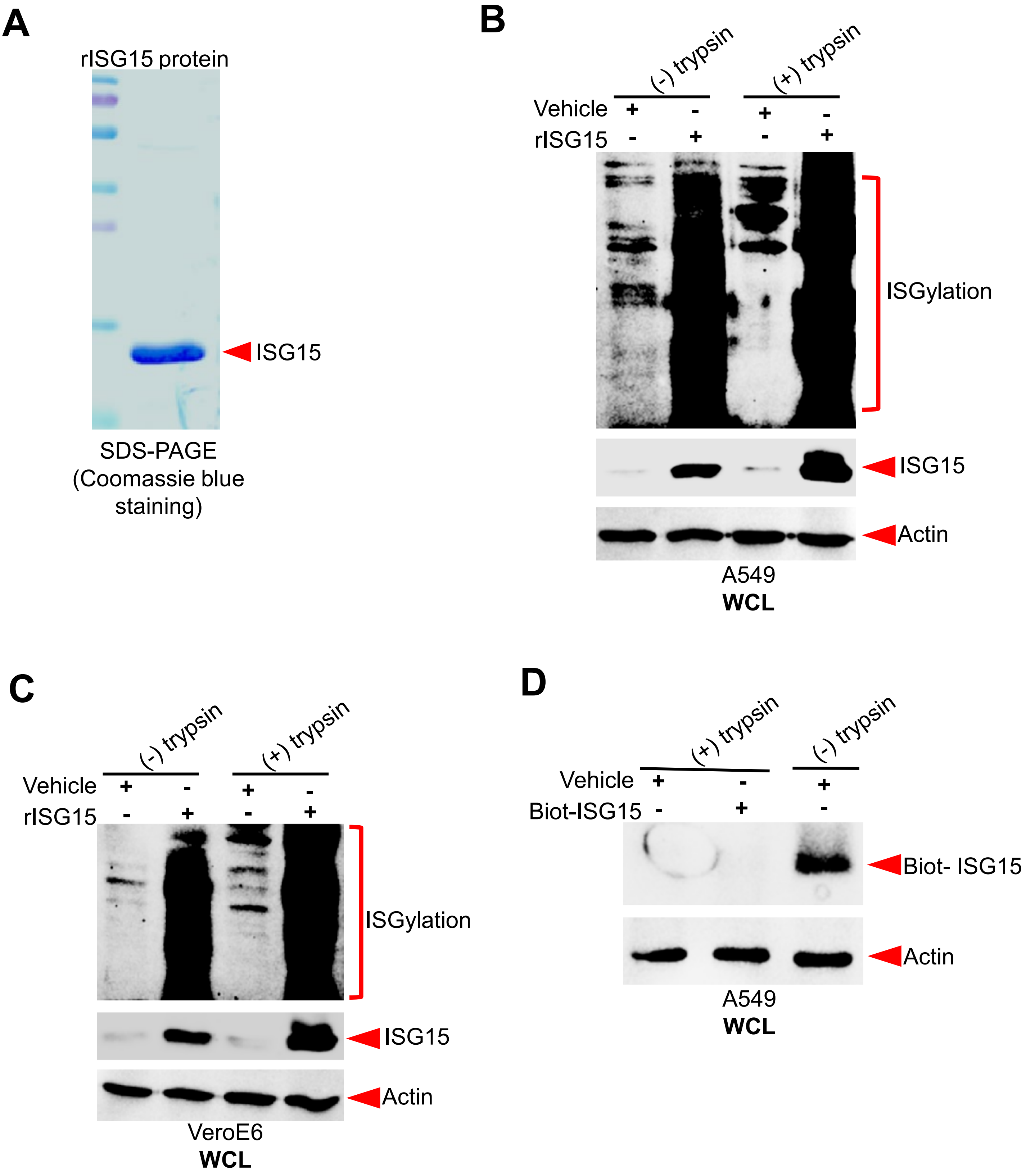

### Supplementary Fig. 3

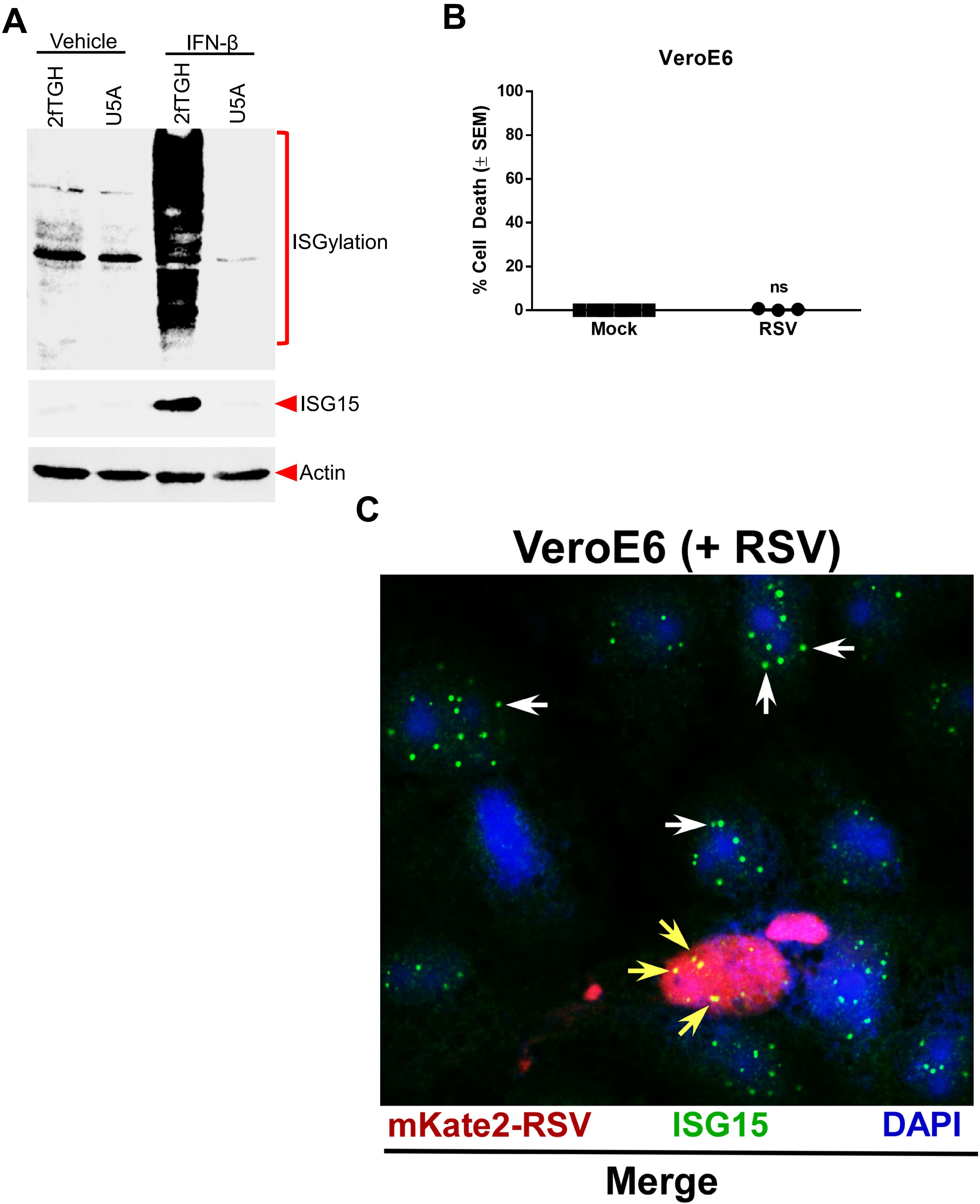

### Supplementary Fig. 4

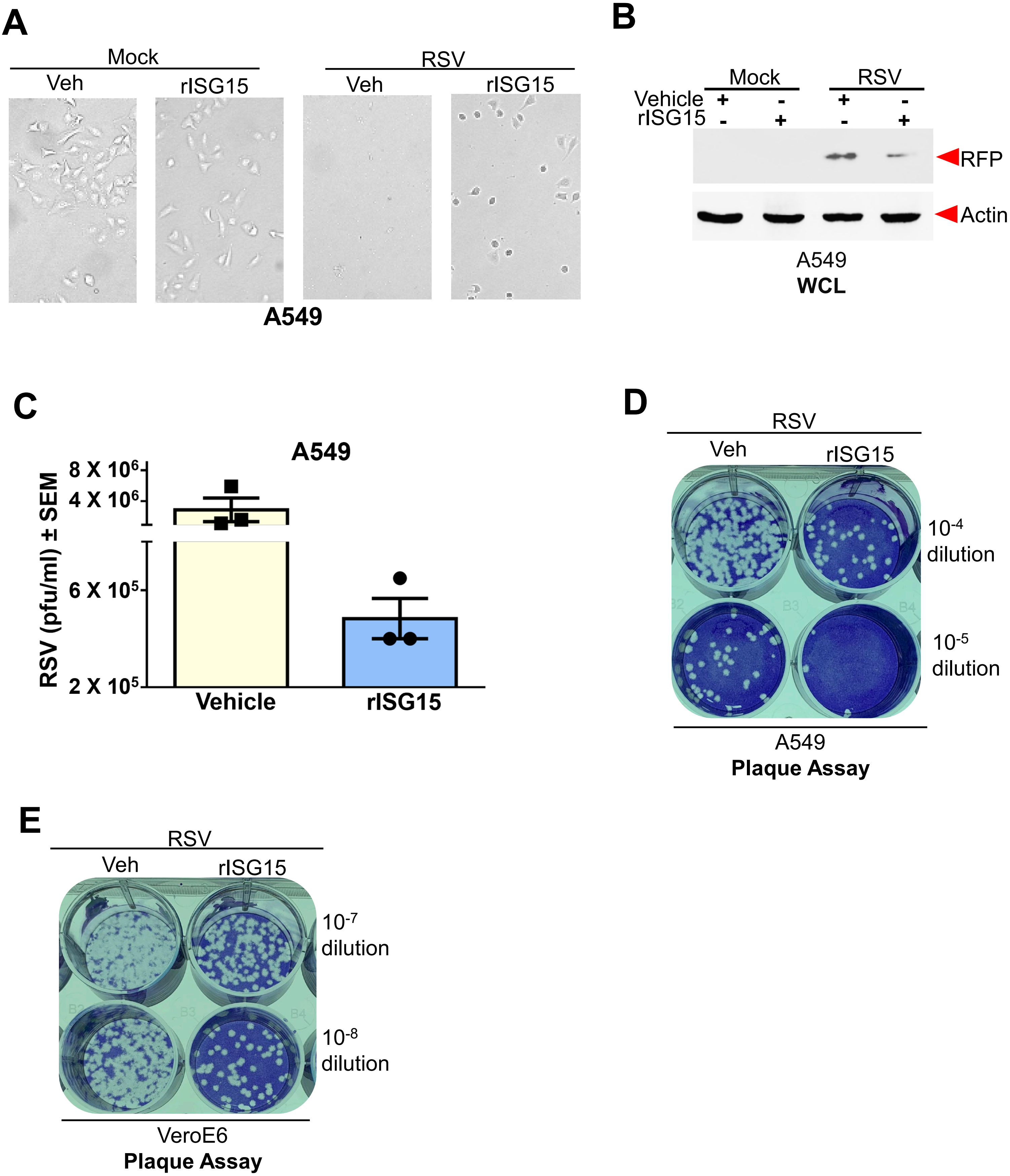

### Supplementary Fig. 5

A

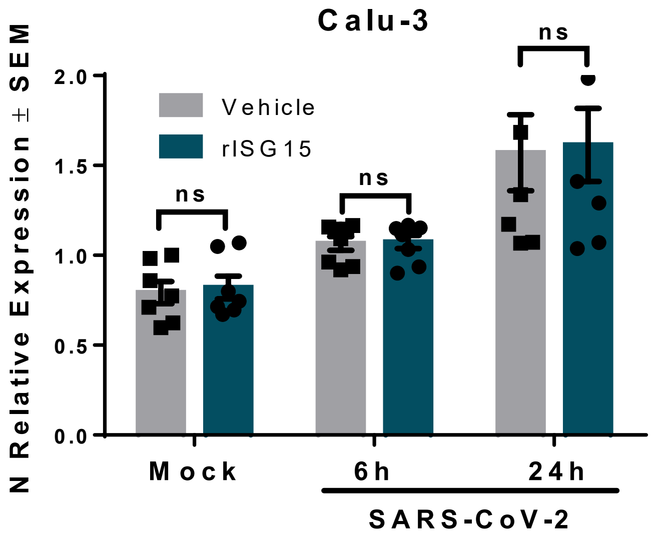


B

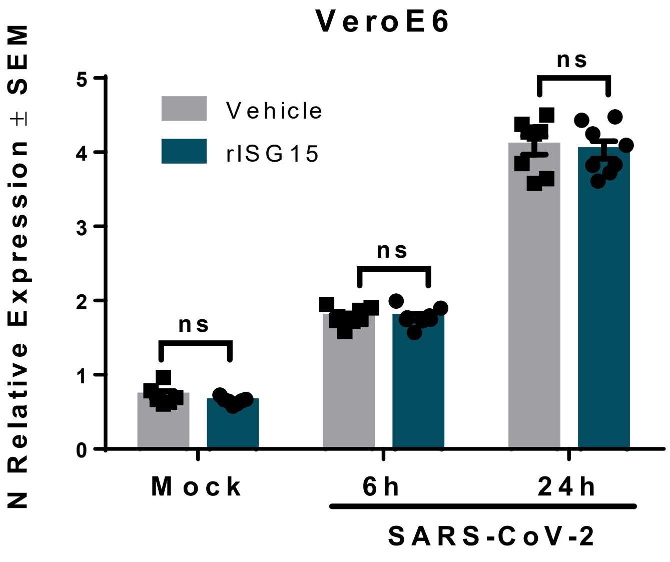

### Supplementary Fig. 6

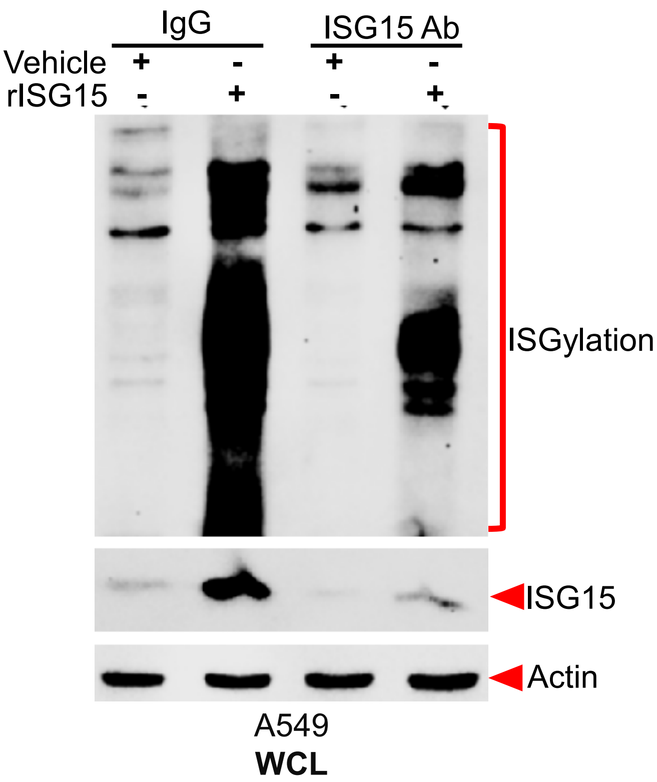

### Supplementary Fig. 7

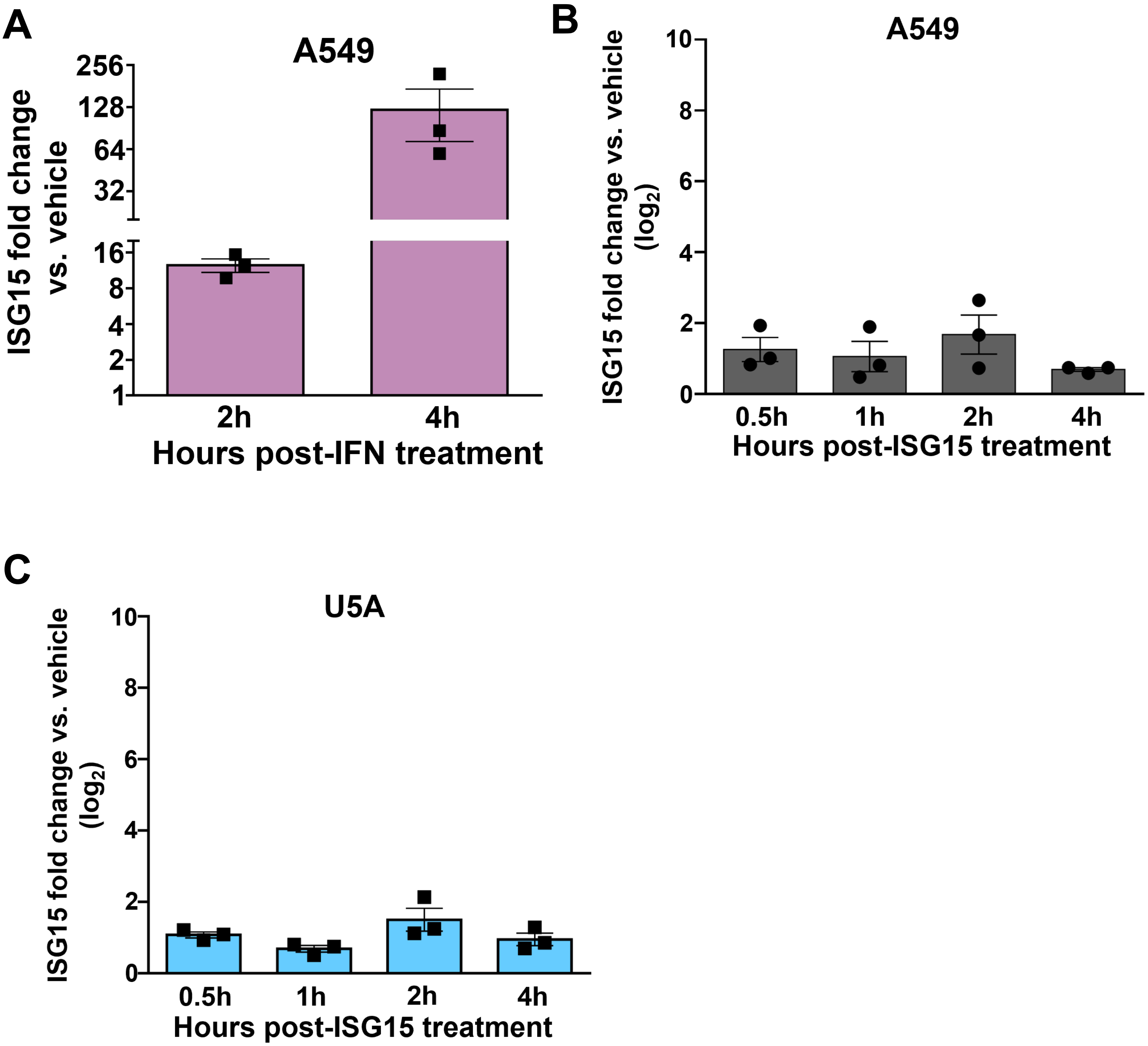

### Supplementary Fig. 8

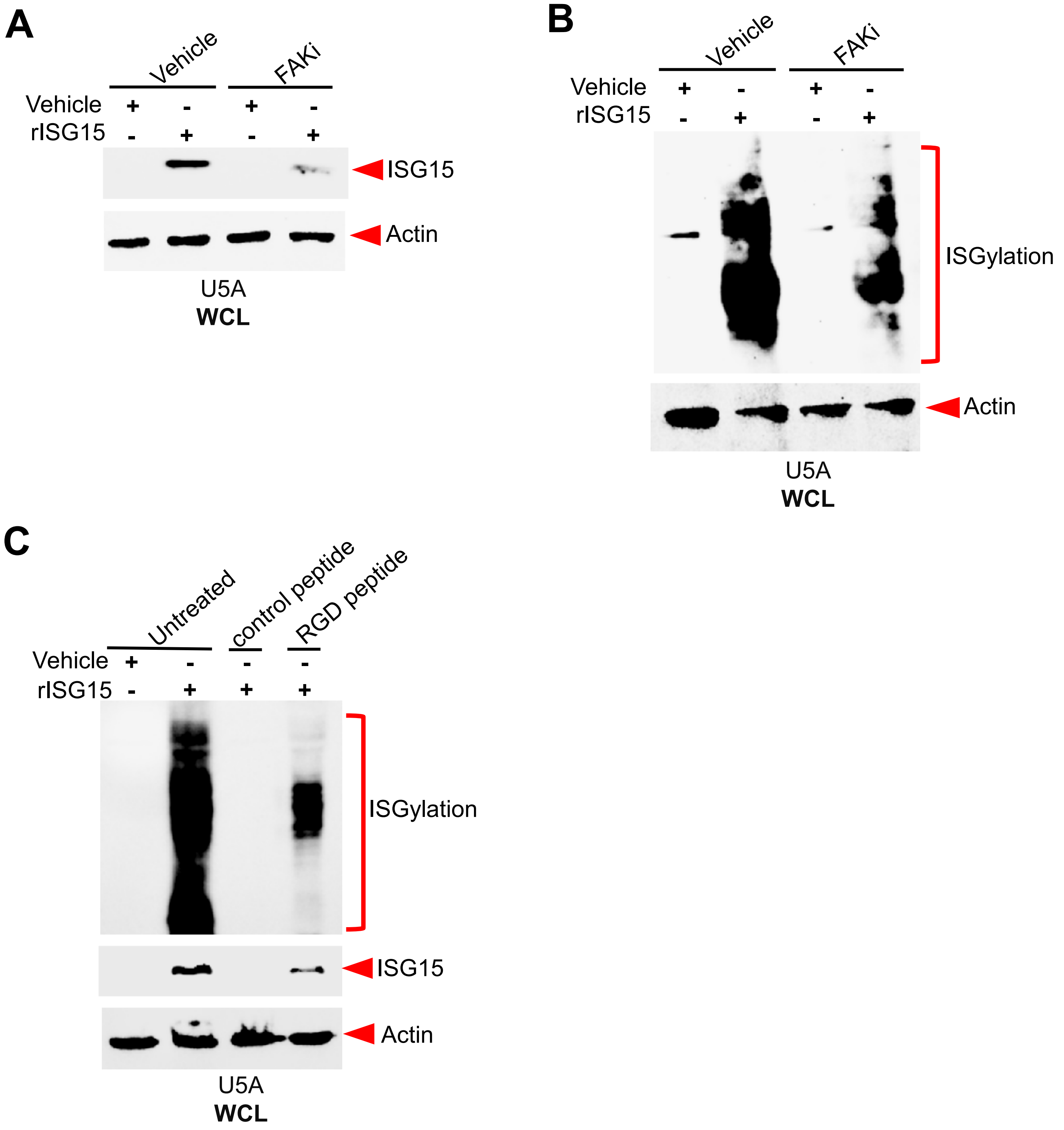

### Supplementary Fig. 9

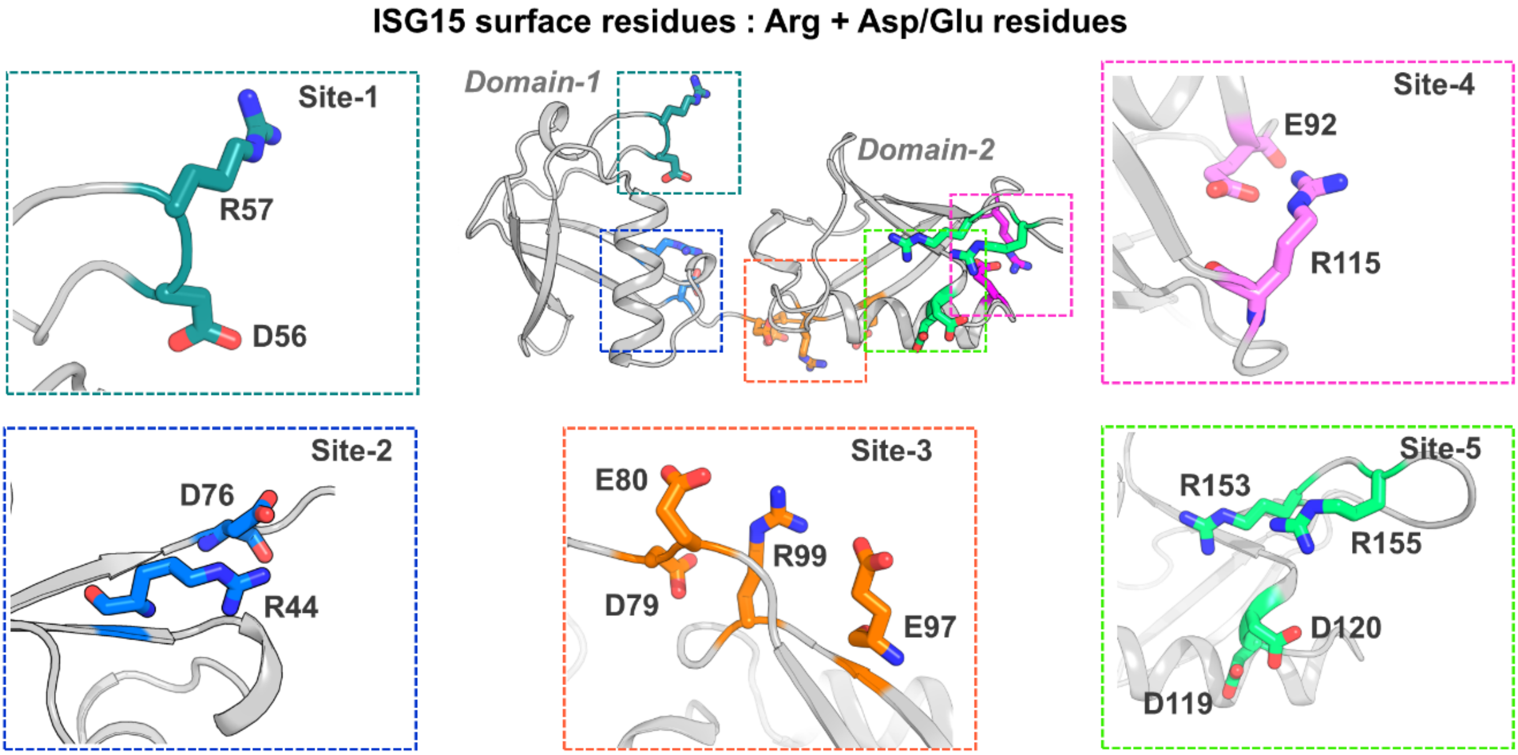

### Supplementary Table

**Table S1.** HADDOCK scores for integrin α5β1 - ISG15 complexes.

| Integrin | ISG15 | score |
|---|---|---|
| RGD motif binding site<br>**(includes α5 and β1 – RGD motif binding residues)** | BLIND<br>**(all residues)** | **-31.0 ± 2.5** |
| | **Site-1**<br>**(Asp56, Arg57)** | **-121.4 ± 5.2** |
| | Site-2<br>**(Arg44, Asp76)** | **-141.8 ± 7.6** |
| | **Site-3**<br>**(Asp79, Glu80, Glu97, Arg99)** | **-165.6 ± 5.3** |
| | Site-4<br>**(Glu92,Arg115)** | **-128.3 ± 4.5** |
| | Site-5<br>**(Asp119, Asp120, Arg153, Arg155)** | **-111.1 ± 2.6** |

#### Supplementary Figure Legends

**Supplementary Figure 1.** A) Human lung epithelial A549 cells were treated with either vehicle control, purified SARS-CoV-2 Spike protein (rSpike, 8nM) or purified S1 subunit of SARS-CoV-2 Spike protein (rS1, 8nM) for 16h. They were also treated with interferon-β (IFN-β) (500 units/ml). WCL was immunoblotted with ISG15 and actin antibodies. B) A549 cells were transfected with either empty vector control plasmid (pcDNA) or RSV Fusion (F) protein plasmid. At 24h post-transfection, washed cells were incubated with fresh media without plasmids for 24h. Whole cell lysate (WCL) was subjected to immunoblotting with ISG15 and actin antibodies.

**Supplementary Figure 2.** A) SDS-PAGE analysis of purified recombinant ISG15 protein (rISG15). A549 (B) and VeroE6 (C) cells were treated with either vehicle control or rISG15 (5μg/mL). At 6h post-treatment, washed cells were either untreated or treated with trypsin before collection of WCL. WCL was immunoblotted with ISG15 and actin antibodies. D) A549 cells were treated with either biotinylated vehicle control (biot-Veh) or biotinylated-rISG15 (biot-ISG15). At 6h post-treatment, washed cells were treated with trypsin before collection of WCL. In another experiment, cells were treated with biot-ISG15 briefly followed by collection of WCL in the absence of trypsin. WCL was subjected to western blotting with avidin-HRP (to detected biotinylated-rISG15) and actin antibody.

**Supplementary Figure 3.** A) 2fTGH (IFN-competent) and U5A (IFN-incompetent) cells were treated with interferon-β (IFN-β, 500U/mL) for 6h before collection of WCL for immunoblotting with ISG15 and actin antibodies. B) Supernatants from mock and RSV infected (48h post-infection) VeroE6 cells were used to quantify cell death by lactate dehydrogenase (LDH) assay. ns; non-significant. C) Magnified confocal merged image from Fig. 4E. VeroE6 cells infected with mKate2-RSV (red) was labelled with ISG15 antibody (green) and DAPI (blue). The merged image of the infected cells shows ISG15 expression in uninfected cells (green dots shown with white arrows) and infected cells (yellow dots shown with yellow arrows).

**Supplementary Figure 4.** A) Bright field microscopy of A549 monolayers treated with vehicle control (Veh) or rISG15 prior to infection with RSV (MOI=1, 16h). B) A549 cells were pre-treated with purified recombinant ISG15 protein (rISG15, 5μg/mL) or vehicle control for 8 hours prior to infection with mKate2-RSV for 48h before collection of WCL for immunoblotting with RFP (to detect mKate2) and actin antibodies. C) Infectious viral titer following RSV (MOI=0.1) infection of A549 cells pre-treated with either vehicle control or rISG15 (5μg/mL). A representative plaque assay showing RSV infectious titer in A549 (D) and VeroE6 (E) cells pre-treated with either vehicle control (Veh) or rISG15 (5μg/mL).

**Supplementary Figure 5.** SARS-CoV-2 infectivity in Calu-3 (A) and VeroE6 (B) cells pre-treated with either vehicle control or rISG15 (5μg/mL). Infectivity was analyzed by performing RT-qPCR for SARS-CoV-2 nucleocapsid (N) protein expression. ns; non-significant.

**Supplementary Figure 6.** Extracellular ISG15 blocking antibody. A549 cells were treated with rISG15 (1μg/mL) in the presence of either control IgG or polyclonal anti-ISG15 antibody (1μg/mL). WCL was immunoblotted with ISG15 and actin antibodies.

**Supplementary Figure 7.** (A) ISG15 mRNA expression in A549 cells treated with interferon-β (IFN) (500 units/ml) was analyzed by performing RT-qPCR. ISG15 mRNA expression in A549 (B) and U5A (C) cells treated with rISG15 (5µg/mL) was analyzed by performing RT-qPCR. Cycle threshold (Ct) values for ISG15-specific primers were normalized to values for GAPDH, and the fold change versus vehicle controls for each time point was calculated (2^-DDCt).

**Supplementary Figure 8.** A) Type I interferon incompetent U5A cells were treated with rISG15 (1µg/mL) in the presence of either vehicle control or FAK inhibitor (FAKi, 20 µM). WCL collected from these cells were subjected to immunoblotting with ISG15 antibody to detect unconjugated monomeric form of ISG15. B) U5A cells were treated with rISG15 (1µg/mL) in the presence of either vehicle control or FAK inhibitor (FAKi, 20 µM). WCL collected from these cells were subjected to immunoblotting with ISG15 antibody to detect ISGylated proteins. C) U5A cells were treated with rISG15 (2µg/mL) in the presence of either control peptide or RGD peptide (200 µM). Cells were also treated with rISG15 in the absence of any peptide (untreated cells). WCL collected from these cells were subjected to immunoblotting with ISG15 and actin antibodies.

**Supplementary Figure 9.** The surface-exposed Arg and Asp or Glu residue combinations on ISG15. ISG15 has five different sites with exposed Arg-Asp/Glu residue combinations on its solvent-exposed surface.
